# Trends in the representation of research on model organisms in scientific literature

**DOI:** 10.64898/2026.03.03.709331

**Authors:** Conrad Fallon, Xinyi Li, Gilberto Alvarez-Canales, Mariam Museridze, Nicolas Gompel

## Abstract

Research using model organisms to tackle questions in life sciences and biomedical sciences has been in the spotlight of scientific literature for the better part of the twentieth century. This attention has perceptibly faded over the last twenty years, at least. We set to document this process by examining the publication trends of 48 journals encompassing a broad range of topics and impact factors for eight classic model organisms. We found that the representation of model-organism research has been in continuous decline in the last three decades, with a significant acceleration since 2010. We investigated the origin of the change, from the size of research communities and the time to publication, to the shifts in topics and in use of model organisms. While model organism communities appear stable, model organism papers are outpaced by the rest of scientific literature. Also, among papers using model organisms, we note a progressive shift toward applied research, with differences between model organism species. The mouse, in particular, remains one of the most widely used mammalian systems for studying human diseases, while non-mouse model organisms continue to be used predominantly to dissect mechanisms of life. We finally examined how funding has evolved in parallel with the fading representation that we measured for the future of life sciences and reflect on the consequences. Fundamentally, model organisms afford a direct access to causality in life sciences and their fading from the picture may impact life sciences as a whole. More pragmatically, it will also affect funding, and thereby jeopardizes the maintenance of model organism resources such as repositories built over decades.

## Introduction

Each scientist wants the fruit of their research to be visible. This is particularly true for studies of fundamental processes of broad relevance, which directly benefit the scientific community, and ultimately society as a whole. Had Mendel’s work not remained largely ignored for over three decades, life sciences would probably look different today. Our contemporary publishing system, where visibility is linked to a journal’s impact factor, is reportedly based on the novelty and relevance of findings. In life sciences, model organism (MO)-based research has become the standard approach during the twentieth century to identify novel processes and elucidate their principles, even if the limit of the approach has been acknowledged (*1*).

Two main reasons have accelerated this transition in deciphering the mechanisms of life. First, the recognition that molecules and fundamental mechanisms are conserved across the tree of life. Second, the possibility to test causality directly in whole organisms through controlled experimental perturbations (including large-scale experiments and genetic manipulation) with limited noise (reduced genetic variation, standardized breeding conditions). Adding to this a historic depth of knowledge with which different phenomena can be connected has made MOs an essential tool for the identification of genes of biomedical interest, the dissection of their cellular and molecular functions, as well as the unraveling of entire processes in which they are involved (*2, 3*). Bibliometric surveys of model MO research clearly reflect this trend over the twentieth century (*4, 5*), with the exponential growth of literature linked to technical developments such as large-scale mutagenesis or transgenesis. The genetics of animal segmentation, the bases of the immune system, or the logic of the olfactory system provide iconic illustrations: worked out in a handful of systems, but relevant to large scientific and medical communities.

Among MO organism communities, however, there is a diffuse and growing impression that it has become much harder to publish significant work in top-ranking journals than it used to be even a decade ago. A recent editorial reaffirming the value of model organisms in scientific thinking and training hinted at the decline in funding and citation of MO-based research (*6*). To substantiate the impression, we set to examine bibliometric data relating to MO-related primary research literature. We compared the publication trends and the representation of research with the main model organisms among scientific journals with different impact factors over the last three decades. Our results highlight an overall steady and alarming decay in the visibility of MO-based research in all journal categories. We explore the possible causes of this evolution, including the size of each MO community, the time required to publication, and the type of research published using MOs, as well as the relationship to research funding, and discuss their consequences on the future of life sciences.

## Results

### Consistent decline in the representation of model organism research across scientific journals

To investigate trends in the publication amount of MO research over the past three decades, we retrieved data from the Web of Science (www.webofscience.com/wos/) on articles published between 1995 and 2024 for eight classical model organisms: *Escherichia coli* (bacteria), *Saccharomyces cerevisiae* (yeast), *Caenorhabditis elegans* (nematode), *Drosophila melanogaster* (fruit fly), *Danio rerio* (zebrafish), *Xenopus tropicalis* (frog), *Mus musculus* (mouse), and *Arabidopsis thaliana* (mustard plant) (see methods for details). The analysis included 48 scientific journals with impact factors ranging from 2.5 to 55. All journals publish primary research articles; some are generalists, while others target specific biological fields (Table S1). We examined how the representation of MO research evolved over time. Specifically, we surveyed the total number of primary research articles published by these outlets, among those, the number of MO-related papers, and finally the proportion of papers published each year that use MOs (MO ratio) (Fig. 1A and S1A).

**Figure 1.**
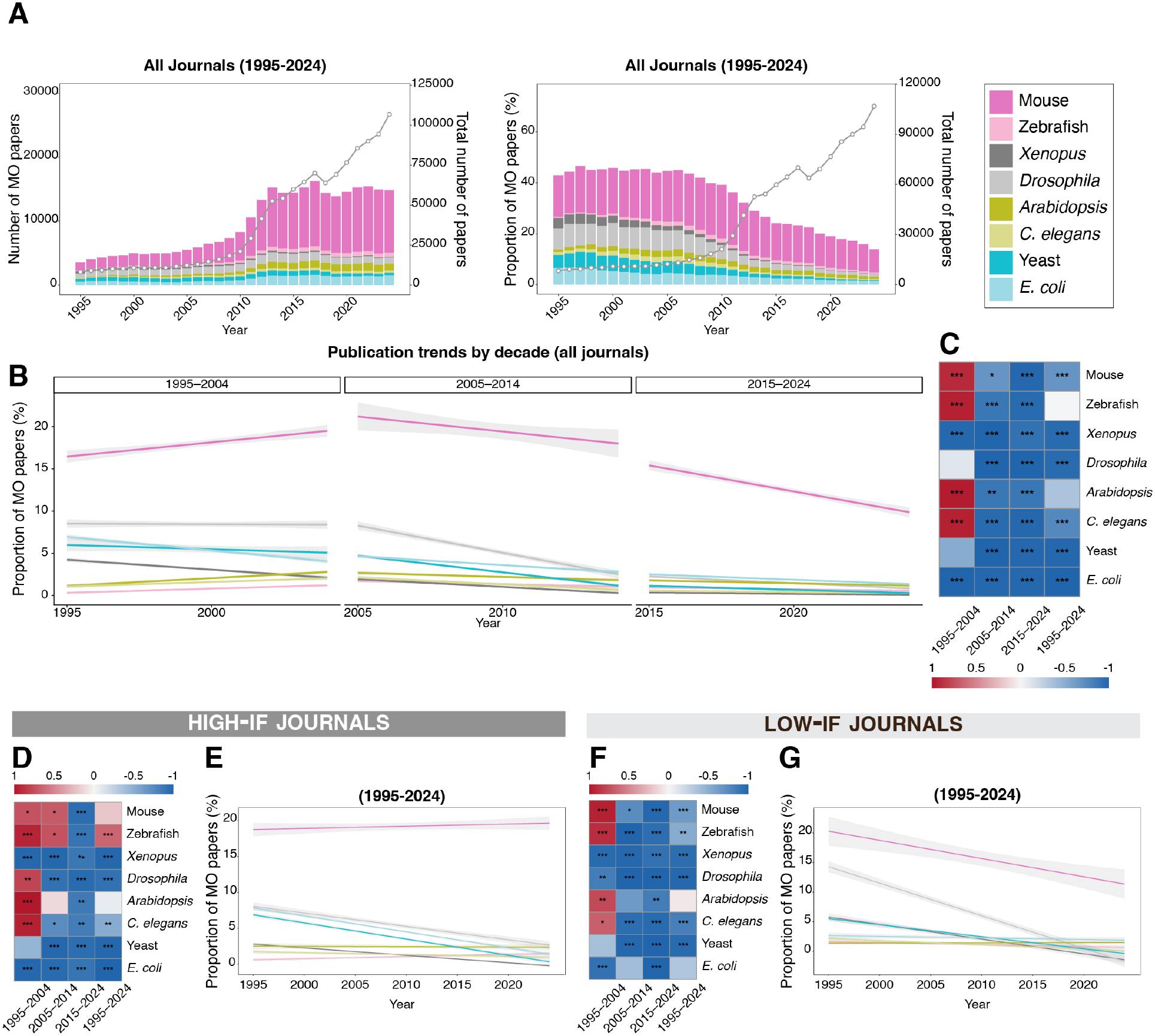
Publication trends of different model organisms in selected scientific journals. **A**. Changes in the number of publications (left) and their proportion as a percentage of all papers (right) for different model organisms across all selected journals listed in Table S1 for the period 1995–2024. Each colored bar represents a specific model organism (color code right to panel applies to the entire figure). The grey line shows the total number of research papers published by the same outlets. **B**. Decade-wise trends in publication proportions across all selected journals. Each panel shows one decade (1995–2004, 2005–2014 and 2015–2024). Linear regression lines with 95% confidence intervals show the trends for each model organism within that decade **C.** Correlation heatmap denotes the temporal trends in model organism publication shares across all selected journals. For each organism and journal, we calculated the correlation (Pearson correlation coefficient, r) between its publication proportion and time across successive decades. The color of each cell indicates the direction of the trend: red for a positive correlation (increasing publication share of that MO over the decade), blue for a negative correlation (decreasing share). The statistical significance of each correlation is separately denoted by asterisks within the cell. **D.** Correlation heatmap of publication trends for model organisms in all high-IF journals, similar to (**C**) **E**. Trends in publication proportions across high-IF journals across the period 1995–2025. Linear regression lines with 95% confidence intervals show decade-specific publication trends for each model organism. **F**. Correlation heatmap of publication trends for model organisms in all low-IF journals, similar to (**D**). **G**. Trends in publication proportions across low-IF journals across the period 1995-2025, similar to (**E**). Statistical significance in all figures is denoted by asterisks: *, p < 0.05; **, p < 0.01; ***p < 0.001. Corresponding statistical data are provided in Tables S5–S7.

We observed a clear declining trend in the MO ratio across all journals. From 1995 to 2024, their relative share decreased by approximately 68%. This is largely explained by the total number of scientific papers increasing exponentially from around 10,000 to over 100,000 per year during this period. Meanwhile, the number of MO papers has increased more slowly, and stagnated in the last decade, resulting in an overall decline in the MO ratio. This trend accelerates particularly around the year 2010, where the number of all papers published rapidly increases, an exponential inflation analyzed by others (*7*). This may be linked to the fragmentation of studies and the foundation of numerous specialized journals, a strategy that serves both academic careers and publisher’s economic incentives (*7*). However, this does not explain why the number of MO papers didn’t increase at a similar rate.

While each MO is affected to different extents by the trend, none escapes it (Fig. S1A), with *Drosophila* taking the highest toll. To quantify and validate this observed reduction, we performed Pearson correlation analyses between publication proportion and publication year for each model organism, both across the full study period (1995–2024) and across successive decades. This allowed us to quantify the strength and direction of the temporal trend and to test whether the trend was consistent across organisms and time windows (Fig. 1B, C). The results showed strong negative correlations for all model organisms across most journals, except for zebrafish and *Arabidopsis*. A more detailed temporal analysis revealed that the decline for mouse, zebrafish, *Arabidopsis* and *C. elegans* began later compared to other organisms (starting in the 2005–2014 period; Fig. 1B, C). For mouse and zebrafish this may be attributed to their stronger alignment with contemporary biomedical priorities—such as drug metabolism studies or disease research (*8–11*).

Most strikingly, in the most recent decade (2015–2024), the proportion of publications of every model organism studied declined significantly. Critically, this decline is also reflected in the changes of absolute number of papers. Except for mouse, zebrafish and *Arabidopsis*, all other model organisms exhibited a decrease in their publication counts over the 2015–2024 period, with *Drosophila, Xenopus* and yeast being most significantly affected (Fig. S2). This finding confirms that the observed trend represents a genuine contraction in the output of MO research, suggesting a broad and concurrent shift away from MO-based research across the publishing landscape. We then sought to understand the origin of these dynamics. We first wondered whether the trend was uniform among journals, or whether it was driven by a particular category, such as high-impact factor journals. We also wondered whether the scientific communities supporting MO research had shrunk over the years, and whether the time needed to publish a paper has changed. Also, as MO have been extensively used to tackle fundamental questions in biology, we examined whether a recent push for more applied research had limited the growth of MO research. Finally, we examined the relationship between research funding and publication rates.

### Comparing high-impact-factor to lower-impact-factor journals

We first asked whether a particular group of journals was driving the global trend that we observed. Our first hypothesis was that this was a trend affecting top-tier journals, where MO articles are outcompeted by emerging research areas (such as AI, cross-disciplinary studies, and organoids, which have seen exponential growth in recent decades, Fig. S3), and lower-impact-factor journals remaining more likely to accept MO papers. We therefore parsed the journals into two groups: high-impact-factor journals (high-IF, impact factor > 7) and lower-impact-factor journals (low-IF, impact factor < 7) (Table S1; Figs. 1D-G and S1B-E). Briefly, the impact factor reflects the average number of citations received by articles published in a journal over a given period (typically the previous two or five years) and roughly indicates the journal’s influence. We designed the groups such that for each high-IF journal, there is a low-IF counterpart. For instance, among generalist journals, we chose *Proceedings of the Royal Society B-Biological Sciences* (IF = 4.4) as a low-IF counterpart of *Nature* (IF = 55).

Surprisingly, our findings reveal that the proportion of publications focused on model organisms declined significantly in both journal categories (Fig. 1D-G and S1B-E). The magnitude of decline was markedly greater in low-IF journals, with a relative reduction of approximately 76% from 1995 to 2024 (Fig. S1D), compared to 39% in high-IF journals (Fig. S1B). Examining the entire period (1995–2024), the declines for mouse, zebrafish, *Drosophila*, and *C. elegans* were more pronounced in low-IF journals (Fig. 1G and S1D). 1995–2004 saw rising representation of several model organisms in both high- and low-IF journals (Fig. S1C, E). The rapid decline in the MO ratio around 2010 is first obvious in low-IF journals, followed a few years later by high-IF journals (Fig. S1B, D).

When comparing individual journals (Fig. S4), the decline is pervasive but not uniform. It may largely be driven by the rapid increase in output by new open-access journals and large-volume multidisciplinary journals, (sometimes referred to as ‘mega journals’) since around 2010, causing the overall trend to reflect their changing publishing profiles (Fig. S5). This is especially noticeable in journals such as *Nature Communications* (43% of all high-IF articles in 2024), *Scientific Reports* and *PLoS One* (together nearly 60% of all low-IF articles in 2024), which each produce over 10,000 articles a year with a relatively low or decreasing proportion of MO papers. This suggests that the global increase in scientific literature is primarily driven by the appearance of new outlets, which may drive part of the global reduction in MO ratio. However, the MO ratio tends to decrease, even among older journals publishing consistent or decreasing numbers of papers, suggesting it is not just the effect of more papers from every field diluting the proportion of MO papers. This appears to be relatively consistent across high-IF and low-IF journals, in different fields and for different MOs (Figs. 1D, F and S6), in spite of heterogeneities among geographical regions where research is done (Fig. S7).

In summary, the decline in MO research is a widespread phenomenon observable across the scientific publishing landscape. It does not appear to be driven by a particular category of journals, and all model organisms, in spite of different rates of decline, are affected in their representation in modern scientific literature. The fact that MO-paper numbers have not increased for a decade suggests they are not able to take advantage of this recent explosion in publishing (*7*) and this is visible in the decreasing MO ratio.

### The decline does not result from shrinking communities or longer time to publication

We next sought to determine whether the decline in the MO ratio stems from an erosion of their respective research communities. The MO ratio could drop because of the output of each lab remaining constant but the number of labs reducing, or the number of labs remaining constant or increasing but their output falling behind average or being diluted by new fields, or both. Getting exact numbers of researchers using each MO is difficult, so as a proxy we used past conference attendance data for annual meetings on MO research: namely, the Yeast Genetics Meeting, the International Worm Meeting, and the Annual *Drosophila* Research Conference, as well as the European Zebrafish Meeting and the European *Drosophila* Research Conference (Figs. 2A-F and S7A-B). The total attendance at the yeast, worm and fly conferences has remained stable over the past 10 years, suggesting the communities may have not changed in size significantly. Attendance at the European Zebrafish Meeting exhibited clear growth from 1999 until its last recorded year (2017), indicating an active and expanding research community up to that point (Fig. S8A). Furthermore, the total attendance at virtual meetings in 2021, in the midst of the COVID pandemic, increased substantially, suggesting that the main barriers to these conference participations are likely geographic distance and financial constraints rather than a lack of research interest or shrinking communities.

**Figure 2.**
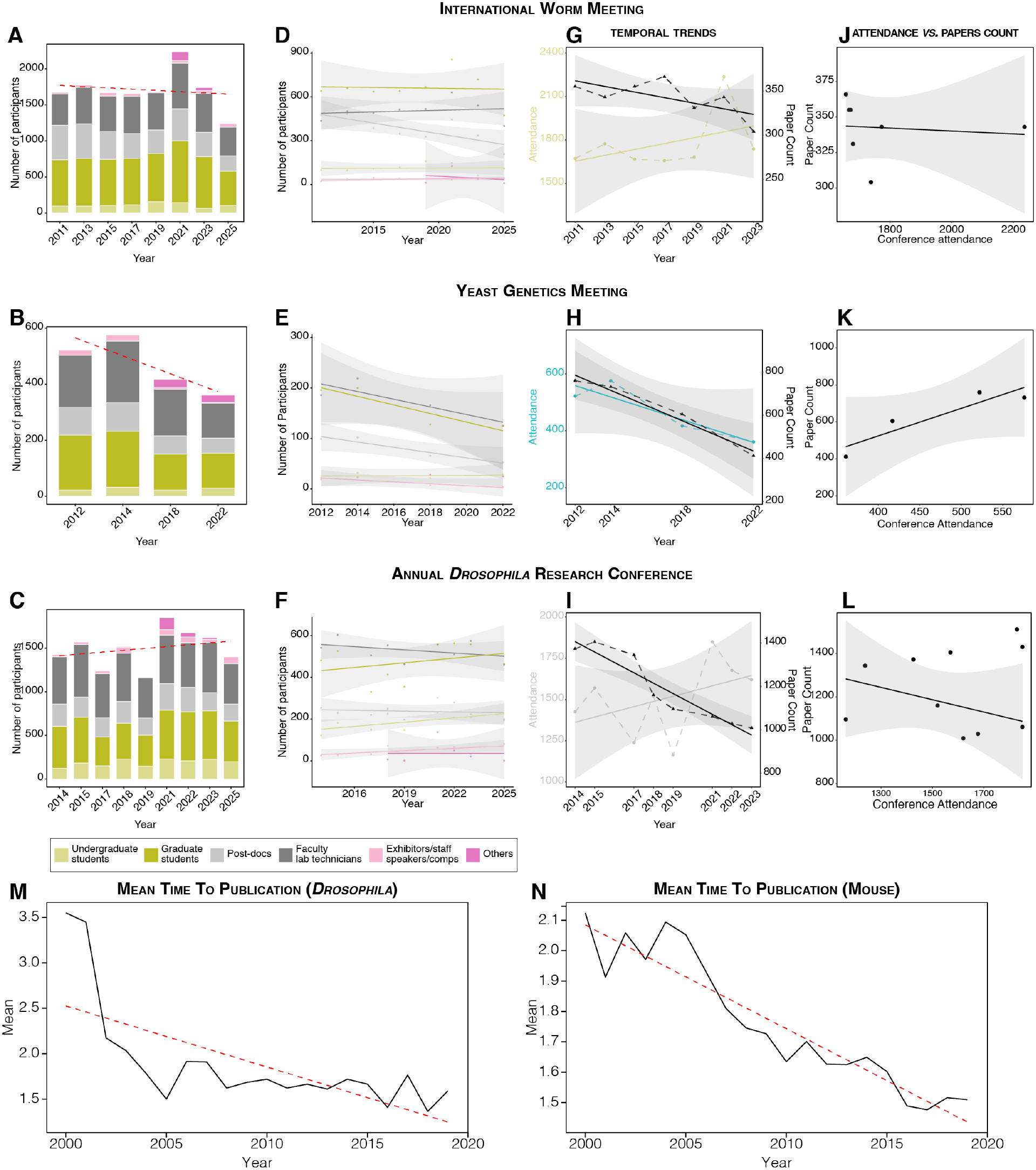
Trends in Model Organisms conference attendance and publication temporal dynamics. **A–C.** Number of participants at international Genetic Society of America (GSA) conferences for *C. elegans* (**A**), yeast (**B**), and *Drosophila* (**C**). Stacked bar plots show participant categories (see inset for color code). Red dashed lines indicate linear trends over time, with Pearson correlation coefficients (r) and p-values as follows: (**A**) *C. elegans*: r = –0.157, p = 0.7105; (**B**) yeast: r = –0.906, p = 0.0936; (**C**) *Drosophila*: r = 0.271, p = 0.4812. **D–F.** Temporal trends in conference attendance by participant category for the same GSA conferences: *C. elegans* (**D**), yeast (**E**), and *Drosophila* (**F**). Each colored line represents one participant category (color code as in **A–C**). Linear regression lines are shown with 95% confidence intervals (shaded areas). Corresponding statistical data are summarized in **Table S8. G–I.** Dual-axis time series plots showing annual changes in conference attendance (colored dashed lines, left y-axis) and publication output changes (black dashed lines, right y-axis) for *C. elegans* (**G**), yeast (**H**), and *Drosophila* (**I**). Solid lines represent linear trends, with shaded areas indicating 95% confidence intervals. **J–L.** Correlation between annual conference attendance and publication count for each model organism. Pearson correlation coefficients (r) and p-values are as follows: (**J**) *C. elegans*: r = –0.11, p = 0.82; (**K**) yeast: r = 0.92, p = 0.08; (**L**) *Drosophila*: r = –0.39, p = 0.33. **M–N.** Temporal trends in mean delay between grant award and first resulting publication. *Drosophila* (**M**): r = - 0.68, p <= 0.001, and mouse (**N**): r= -0.94, p<0.0012. Dotted red lines are linear regression lines.

In summary, among the four model organisms for which we obtained core conference attendance data, three showed relatively stable attendance over the past decade, indicating that the size of their active research communities has not significantly contracted. To further test whether community size could explain the decline in research output, we compared yearly publication output with same-year conference attendance (Figs. 2G-L and S8C-F). If community size were the primary driver of output, we would expect a positive correlation. We found a positive correlation for yeast and zebrafish, but a negative correlation for *Drosophila* and *C. elegans*, with most correlations not reaching statistical significance. Thus, regardless of the direction of the relationship, community size does not appear sufficient to explain the overall decline in MO publication representation. If community size is not the cause, one possibility is a shift in the time to prepare a scientific publication, as a result of higher quality standards, demands on depth, etc. To test this possibility, we used data from NIH Reporter (https://reporter.nih.gov/) to measure the evolution of the time between funding acquisition and first resulting publication for individual *Drosophila* and mouse projects. Specifically, we measured the average delay between when a grant was awarded and its first resulting publication. Contradicting our expectation, the mean delay between funding acquisition and first resulting paper did not increase (Figs. 2M-N and S8G-N). Rather, we observed a statistically significant negative trend in both *Drosophila* and mouse. As more recently awarded grants will have fewer publications, they create a possible bias. We therefore also analyzed works published within a 5-year window after start of all grants. Consistently, this more controlled approach also shows negative trends, although it loses the significance (Fig. S8G–N). These observations suggest that most papers are now published closer to a project start date and may be explained by smaller, more focused papers, possibly reflecting the urge to publish.

Taken together, changes in community size and in the time required to publication do not adequately explain the declining proportion of MO papers. This phenomenon is more likely explained by the rapid development of other research fields, whose growth has outpaced that of traditional MO research. This raises two questions: why is MO research not expanding at a comparable rate, and what types of research are emerging to fill the gap?

### Identifying research foci underlying MO publication trends

The persistent lag in MO-related publications prompted us to examine whether the research environment had shifted away from MO use in recent years. Particularly, we considered whether a shift towards applied research over fundamental research—for which MOs are especially useful—could have driven the decay in the proportion of MO publications.

To track emerging trends in types of published research, we used a large language model (LLM, zero-shot classifier (*12*)) to categorize research articles based on their primary focus: ‘disease research’, ‘applied research’, ‘basic research’, or ‘mechanisms of biological processes’ (Figs. 3 and S9, Table S2). To reduce computation time, we limited our analysis to the high-IF journals. We found the proportion of disease and applied research papers nearly doubled between 1995 and 2024, while the number of mechanistic papers decreased by about a third, and that of basic research papers stayed roughly the same (Fig. 3A). Coinciding with the reduction in the MO ratio (Fig. S1B), the period when published research tends to become increasingly applied and disease-focused occurs between 2005–2014 (Fig. S9J).

**Figure 3.**
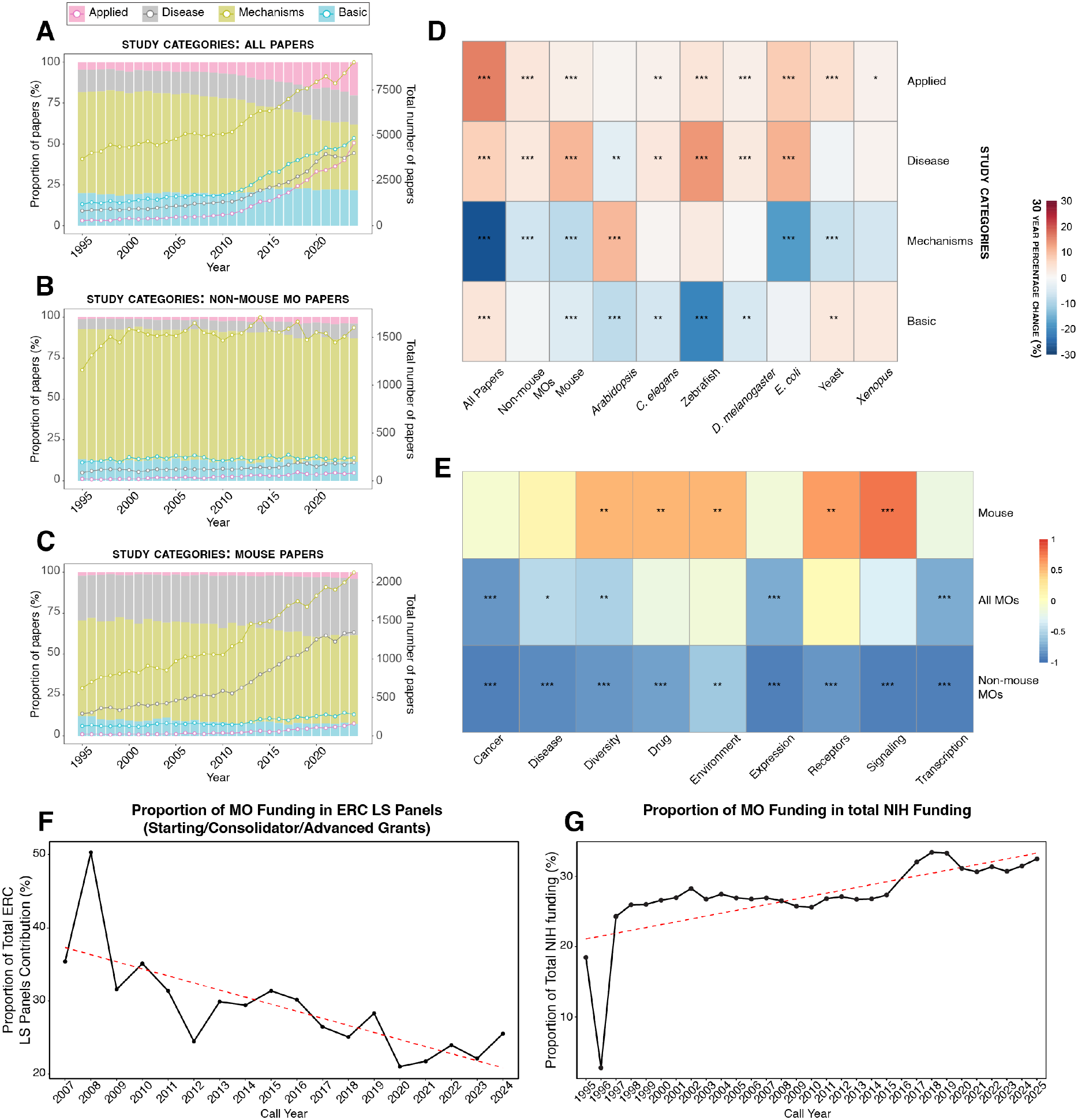
Type of research papers and specific keyword trends in high-IF journals. **A**. All papers of the categories ‘basic’, ‘mechanistic’, ‘applied’ and ‘disease’ according to an LLM classifier based on abstracts published per year from 1995–2024 (left axis, proportion of papers for each year; right axis, number of papers. **B**. Same as (**A**) showing trends for non-mouse MO papers. **C**. Same as (**A**) showing trends for mouse papers. **D**. Heat map of absolute change in proportion of each category over 30 years for all papers, non-mouse model organisms, and each model organism separately. A linear regression of each proportion over time is indicated by the color—corresponding to the change in proportion over 30 years, with the coefficient p-values indicated by asterisks. **E**. Correlation heatmaps for the evolution of the proportion of papers that include the selected keyword (cancer, disease, diversity, drug, environment, expression, receptors, signaling, or transcription) in high-IF journals from 2000–2024. Pearson correlation coefficients (r) were estimated for each of the specific keywords for papers including all MOs, mouse only, and non-mouse-MO research. Blue indicates decreasing trends (r < 0), white indicates no correlation, and red indicates increasing trends (r > 0), as shown on the colormap. Statistical significance is indicated by asterisks within each cell. **F**. Temporal trends in the proportion of ERC funding supporting MO research across LS panels. Pearson correlation coefficient: r = -0.75, p-value: p<0.001. **G**. Temporal trends in the proportion of NIH funding for MO research. Pearson correlation coefficient: r = 0.661, p-value: p = <0.001.

We next investigated whether the proportion of papers published for different MOs could be linked to the change in their representation in high-IF journals (Fig. 1E). We divided our analysis into mouse and non-mouse MOs, since mouse papers represent the majority of MO papers published and have experienced less change in representation over time. We found that while the proportion of non-mouse-MO papers in high-IF journals dropped by around half, their research profile has remained consistently mechanistic (Figs. 3B and S9I-J). This suggests the representation of non-mouse MOs in high-IF journals remains linked to the overall proportion of mechanistic papers published, which has been declining for years. Furthermore, while the number of mechanistic pa-pers using non-mouse MOs has stagnated (Figs. 3B and S9I), the total number of all mechanistic papers has increased (Figs. 3A and S9J), suggesting mechanistic research is increasingly using alternatives to non-mouse MOs. Breaking down each MO separately, most non-mouse MOs have had similar research profiles for the past three decades (Fig. S9I). Notably, however, *E. coli* has increased its disease and applied research proportion *without* changing the total number of papers published (Figs. S2 and S9) while, *Arabidopsis* has become increasingly mechanistic while *in-creasing* in total number of papers published (Figs. S2 and S9).

By contrast, mouse papers have been better able to keep pace with publication trends as a whole; the proportion of mouse papers published in high-IF journals has remained relatively stable over 30 years (Fig. 1E). We found mouse papers tend to be much more disease-focused than other MO papers (Figs. 3B-C and S9F), and have become increasingly so over 30 years; disease-focused mouse research increased from 26% of all mouse papers between 1995–2000 to 33% between 2020–2024. Furthermore, unlike for non-mouse MOs, the total number of mechanistic papers using mice has increased. This suggests two things: that mouse can rely on its use in disease research to maintain a high number of publications as publishing focuses more on disease and applied research; and that mechanistic studies are increasingly using mice over non-mouse MOs. This second point may be linked to the overall increase in disease and applied research, as mice are more closely related to humans, and identifying mechanisms in mice can be more easily applied to human disease studies. These two factors likely enable mouse papers to stay relevant and continue growing.

Finally, to perform a fine-grained inspection of MO use in specific fields of research, we analyzed the frequency of different specific keywords in MO and non-MO papers to link them to changing trends in MO publishing. Furthermore, we explored whether there were associations with applied or fundamental research, and whether this trend remained negative when mouse research was included, compared with the rest of the model organisms.

For applied research we used ’disease’, ’drug’, ’environment’, ’cancer’ and ’diversity’, and for fundamental research we used ’expression’, ’receptors’, ’transcription’ and ’signaling’. The rationale for our selection was that these keywords appeared frequently enough to produce reliable trends in publication for MO and non-MO papers, while still being specific enough to separate applied from fundamental studies (even though some overlap is likely).

We performed Pearson correlation tests to measure how the frequency of selected keywords in MO papers changed relative to the frequency of the same keywords among all papers over time (2000–2024 and 2015-2024). We observed overall negative trends for most keywords across both high- and low-IF journals (Figs. 3E and S10), indicating that MO research constitutes a diminishing portion of all research on the selected topics. Moreover, these negative trends are more pronounced in the correlation and significance scores when mouse research is excluded from the MO tested. This may be explained by mouse still being a prevalent model organism for some of these topics, as revealed when examining the same trends only for mouse papers in high-IF journals (Figs. 3C and S10C). This also aligns with our previous observations that mouse research is not decreasing overall, at least for high-IF journals. Overall, these results support the idea that the negative trend in MO publications is not driven by any specific topic, but rather uniform across fundamental and applied research, with some exception for mouse research.

### Funding trends for model organism research differ between major funding agencies

As a final step to see how MO publishing relates to MO research funding, we performed an analysis of funding data from two large public funding bodies, ERC (Fig 3F, S11-S14) in Europe and NIH (Fig 3G, S15) in the US. The two bodies show contrasting trends in funding for MO research. Concomitant with the decline in MO publishing proportions, we saw a reduction in the total amount of research budget allocated to MOs from ERC, with the total budget spent on MOs dropping from nearly 35% to slightly over 25% between 2007 and 2025 (Fig. 3F). This trend was replicated in half of MO species including *Drosophila* and yeast (Fig S11). Overall suggesting a reduction in relative funding at the same time as MOs become less prominent in the publishing landscape. However, the NIH budget showed the opposite trend. The proportion of NIH funding spent on MOs increased slowly but steadily from around 20% to 30% (Fig. 3G). The absolute amount of funding from ERC decreased slightly between 2013–2023 (Fig S13), but the amount for MOs from NIH increased roughly in line with its increasing budget (adjusted for inflation using the Biomedical Research and Development Price Index) (Fig S15). However much of the increase over 30 years is attributable to a doubling of the funding between 1995 and 2005, which has slowed to a roughly 10% increase between 2005 and 2025. We also analyzed the relation between funding amount and publication rates. Here, in most species, we found a strong positive relationship between the amount of funding received by an MO community each year and the number of papers published related to that MO, strongly indicating that publication and funding are related and that one influences the other (Fig S14, S15). While this was expected, it reaffirms that publishing is a reasonable proxy for the amount of funding a field receives. Overall, these results suggest that any changes to the funding of MO research are likely to be diverse between different funding bodies, and hint at a complex relationship between funding and publishing.

## Discussion

### Is MO research no longer needed?

Our data substantiate the growing impression that MO research is progressively losing momentum in the global landscape of scientific literature. This decline coincides with a notable shift toward applied and disease-focused research topics, raising a natural question: have model organisms already fulfilled their duty in that respect? We argue the answer is no: Model organisms remain indispensable for understanding biological causality, for driving technological innovation, and for sustaining the connection between diverse areas of biology.

Over the past three decades, transformative biotechnologies have repeatedly depended on insights derived from model organisms. The elucidation of RNA interference in *C. elegans* provided a conceptual and technical breakthrough for gene silencing (*13*). Similarly, CRISPR-based genome editing, rooted in bacterial immunity, was rapidly adapted and optimized in established model systems. *Drosophila* remains a powerful *in vivo* platform for testing Cas variants (*14*) or for personalized medicine (*15*), while the zebrafish has enabled large-scale genetic analyses and disease modeling in a vertebrate context (*16*). Likewise, the combination of live fluorescent imaging, optogenetics and connectomics implemented in *Drosophila*, *C. elegans* or mice, has revolutionized our understanding of the nervous system, from the function of single neurons to emerging properties such as learning and memory or spatial navigation (*17–23*). The definition of MOs also affords the study of fundamental cellular processes, such as transcription and translation, in a physiological context, and through a vast range of temporal and spatial scales (*24–27*). In each case, these models bridged the gap between molecular insights and physiological relevance, accelerating both tool refinement and functional discovery.

Foundational insights in experimental biology have often depended on the depth of analysis achievable in well-established model systems. Yet, one might argue that the growing understanding of diverse organisms could diminish the need for classical model systems. Advances in transgenesis, genome editing, and genomics have empowered access to causality in countless species for the better (*28*). This diversification is welcome and broadens our perspective on biological variation with improved phylogenetic coverage. Furthermore, the dialogue between model and non-model organisms can accelerate the rate of discoveries for specific questions. For example, knowledge from *Drosophila* patterning, having been established in the crustacean *Parhyale hawaiensis*, allows the species to be used to understand general principles of regeneration (filling the gap in the phylogenetic tree of regeneration left by *Drosophila*) (*29*). Emerging systems should, however, complement rather than replace classical models. New organisms often lack the decades of accumulated genetic tools, curated mutant collections, stock centers, and community infrastructure that make established MOs uniquely powerful (*30*). These resources cannot be easily recreated. Therefore, while we fully support the development of new models to capture broader biological diversity, sustained in-depth analysis in a limited number of well-supported systems will remain essential for advances across many fields of biology.

Our arguments do not imply that every classical model is indispensable for every biological question. Scientific priorities evolve and certain systems may become less central for particular problems, and subject to what is considered relevant by the research community and funding agencies. The latter depends on many factors such as governmental goals, current world challenges, journal trends in citations, the academic community, historical biases, etc. Nevertheless, the accumulated resources and collective knowledge embedded in established model organisms continue to underpin advances across multiple domains. Model organisms have not exhausted their relevance; they remain essential platforms for discovering and testing the mechanisms that structure living systems.

### Is a proportionate reduction such a bad thing?

Our data shows that rather than a strict decline in the number of MO papers, what has changed in the last decades is the proportion of MO papers in the total number of among all published papers. One may argue that as long as the total number of published MO papers remains steady, at a time when new fields continuously emerge, there is no point worrying about a proportionate reduction in MO publishing. However, given the importance of model organisms to the research ecosystem, their sustained decline in share carries structural consequences—affecting how the field is seen, evaluated, and supported in a publication system built on visibility and metrics. First, declining proportional representation reduces visibility. Unless researchers know specifically what they are looking for, they are generally reliant on journals—or intermediaries—to present papers that may be worth reading. We have shown that these media increasingly draw from a pool that includes proportionately fewer MO papers. Unless they actively seek them out—and short of a monkish devotion to the entire scientific corpus—this means researchers will be less likely to see and cite new MO papers. This is compounded by the recent explosion in articles published in extremely high-volume open-access journals (*7*), whose editorial decisions strongly shape the overall publishing landscape (Fig. S5). Over time, it not only slows the pace of discovery by limiting what scientists read and build upon, it also shapes what is amplified through commentaries, outreach articles, and broader science communication. As a result, MO research is progressively edged out of public and policy discourse.

Consequently, the reduced proportion will make it harder to acquire funding for MO research. For better or worse, securing research funding depends on a track record of publishing—particularly in high-IF journals—and having a high h-index, and is essential to keep a field alive. Despite guidance from funding bodies discouraging the use of such metrics for allocating funds (*31–33*), and thousands of scientists calling for a different model of research assessment (*7, 34*), they remain frequently used to determine hiring decisions (*35, 36*) and correlate with the amount of funding received by labs (*37, 38*). A drop in MO-research funding is already a reality, at least in Europe, even if we don’t show that it is a cause rather than a consequence of MO’s reduced representation in the literature. Further loss of funding for a research field beyond a reduction in publications jeopardizes research infrastructure, which makes the process of doing research harder (*39*). This has the potential to develop into a negative feedback loop whereby a field falling out of favor will find it harder to recover its track record of publishing and struggle to regain its influence. Our analysis shows we may already be seeing this risk at work, as the absolute number of MO papers across all journals has flatlined in the last 10 years and for some species has decreased. In high-IF journals between the periods 2010–2014 and 2020–2024 the number of *C. elegans* papers dropped by 10%, *Drosophila* by 15%, *S. cerevisiae* by 26% and *Xenopus* by 35%, It is not inconceivable that they could eventually become relatively niche species, and other MOs could follow suit.

### Are production-side dynamics driving the decline?

While the visibility feedback loop likely contributes to the sustained proportional decline we observe, it does not fully account for the trend. Additional forces—particularly those operating on the production side of science—may also be driving the shift. These include MO community size and changes in required time for a publication. Contrary to our expectations, neither of these production-side factors appears to explain the decline in the MO publication proportion.

An alternative interpretation is that mouse and *Drosophila* research is becoming fragmented: rather than reducing output or shifting toward longer, larger papers, labs may be producing smaller, more focused papers at a comparable or even faster rate. This trend may be driven by two complementary forces. First, early-career scientists face strict temporal constraints (*40*) and must establish a publication record within limited time frames and under intense competition; their decisions are therefore strongly shaped by the need to secure faster returns (*7*). Second, current editorial review processes may be more strongly influenced by author and manuscript characteristics—such as institutional prestige, team size, and research topic (*41*)—meaning that whether a paper is successfully published may depend on factors beyond the quality of the research itself. In this context, the observation that mouse and *Drosophila* papers are published faster is not surprising: publication outcomes are not determined by research quality alone, but are substantially influenced by the external characteristics described above.

However, if MO research increasingly produces smaller papers, it may offer fewer opportunities for the kind of high-impact, career-defining publications that are prioritized in hiring and promotion. Meanwhile, some expanding areas may offer clearer pathways to high-impact outputs, as indicated by the nearly six-fold variation in acceptance rates across research topics at elite general science journals (*41*). In this context, the opportunity cost of pursuing MO research within current evaluation systems is rising. This may, in turn, influence the decisions of young researchers when choosing their research directions—though further work is needed to test this hypothesis.

Overall, while production-side changes do not appear to explain the decline in MO publication proportion, our data hint at a shift in the nature of MO-research output—toward smaller, more fragmented papers—as a potential contributing factor. This pattern may have implications for the long-term vitality of the field.

### Applied and basic research, a balancing act

The decline in MO publishing may not be an isolated phenomenon, but part of a broader structural shift in the research ecosystem. Our brief analysis shows an increase in the relative number of applied and disease-focused research at the expense of basic research. If we understand applied research as being the consequence of years of basic research being put to practical use, we could consider this a success. The development of mRNA vaccines, ways to design new proteins and drugs, and our ability to understand and react to climate change being notable examples. However, since applied research is built on basic research, a relative reduction in overall basic research could ultimately slow the pace of new discoveries, both practical and fundamental. It is too early to say if the rate of new papers will slow (which seems unlikely at least in the short term), but seeing basic research as an investment in the future and applied research as ‘cashing-in’ knowledge gained, it would be wise to maintain a healthy balance between the two, which means maintaining investment in basic research and model organisms.

While we suspected an effort by funding bodies to promote applied research, funding strategy documents usually encourage a mix of basic and applied research and global data breaking down the amount of funding for each type is not readily available. Our brief analysis of ERC and NIH funding for MOs showed a mixed picture, where the ERC data suggests a trend away from MOs and NIH in favor of MOs, which may reflect a remit to promote applied or fundamental research by each body respectively. In line with this, our analysis of the ERC data showed a shift towards disease and environmental research funding (suggesting a shift towards applied research), and a previous study showed 87% of NIH funding on drug-related subjects in the decade 2010–2019 focused on the molecular targets of drugs more than development of the drugs themselves, suggesting a more basic research approach (*42*). However, whether reduced visibility of MOs itself affects funding is difficult to say; both bodies have access to the same literature but each may have different incentives to promote MO research. One would need to compare the reduction in MO publishing to global MO funding, in a landscape with multiple funding bodies per country, not all of which have easily accessible databases.

Specific cases are indicative of the trend towards applied and non-MO research. For example, a recent announcement by the NIH in the US to prioritize human-focused research, includes a rule that funding for animal model systems must “support human-focused approaches such as clinical trials, real world data, or new approach methods” (*43*). Additionally, private research funding likely influences the shift towards applied and non-MO research. A European Commission report (*44*) shows funding is increasingly drawn from private (specifically business and enterprise) sources, which typically favor applied research. In the EU, US and China, the proportion of R&D funding from public sources significantly dropped between the year 2000 (EU 36%, US 26%, China 34%), and 2022 (EU 32%, US 18%, China 18%), with a particular drop in the US and EU starting around 2010. This shift may even motivate specific bodies like NIH to promote MO research to compensate, but in some cases public funding is itself becoming more applied. In the United States, the federal government has reallocated its portfolio: between 2000 and 2017, the share of funding for basic research contracted from 58% to 42%, while applied research expanded from 27% to 35% of the total. Thus, within public funding, the balance is shifting towards application (*45*). Together, the reduced proportion of MO publications may be the result of a general shift towards private research funding and increased applied research within public funding bodies. An important unanswered question of this research is how uniform the trend is among funding bodies globally and how they influence and are influenced by the rate of MO publication.

### Concluding remarks

Taken together, our findings reveal a stark drop in the proportion of MO-based papers across all types of scientific journals publishing studies in life sciences. This is in conjunction with an apparent refocusing of research away from generalizable and fundamental studies toward applied research. The long-term impact of MO-research discoveries is a blind spot of our study and is generally difficult to quantify. While journal impact factors reveal short-term trends, the heuristic impact on the general research community takes time to become clear. For these reasons, the decline of MO-research representation should at the very least be a wake-up call, within model organism communities. It should also draw the attention of journals and funding agencies to how their decisions may affect MOs and basic research. MOs continue to provide an incalculable benefit to research as a whole, but will always rely on the active support of decision makers, and from their community to justify their existence. One solution may be to refocus MO research to include more applied and disease focused fields as may have worked for mice. Another may be to redress the imbalance of public-private funding or incentivize basic research funding to ensure the continued investment in and visibility of basic research. In any case, the declining proportion of MO research is not an inevitable fate but a signal—one that calls for both self-examination within the MO communities and a broader conversation about how we evaluate and support the full diversity of scientific approaches that together advance our understanding of life.

## Materials and Methods

### Keyword-Based Retrieval for MO Publication Database

To analyze the publication trends of MO research, we have selected 48 journals with high impact factors (IF>7) and lower impact factors (IF<7), either generalists, or specialized in a domain of life sciences (Table S1). We then applied filters of the Web of Science (www.webofscience.com) to retrieve references with the keyword “*Drosophila*”, OR “zebrafish” OR “*Xenopus*” OR “*Escherichia coli*” OR “*C. elegans*” OR “*Saccharomyces cerevisiae*” or “*Arabidopsis*” in either of the following fields: Title, Abstract, Author Keywords (input by authors) or Keywords plus (automatically generated by Web of Science). We limited the search to selected journals listed in Table S1 for the years 1995–2024. We also limited the search to the category “Article”, removed editors’ comments, letters, etc., but kept proceeding papers and retracted publications (generally <0.1%). This retrieval produced a dataset comprising annual publication counts for each model organism in each journal, along with the total number of articles published per journal per year. These data enabled the generation of stacked bar plots, which visually distinguish the contribution of each model organism to the total publication output and clearly illustrate how these contributions have shifted over the 1995–2024 period.

### Statistical Analysis of Publication Trends

Publication data were retrieved from the Web of Science for eight model organisms across a selection of high- and low-impact factor journals spanning 1995–2024. Annual publication counts and total articles per journal were extracted and compiled into structured datasets. Years without publications were not included in subsequent analyses. For each journal and year, the proportion of papers focusing on a given model organism was calculated as : 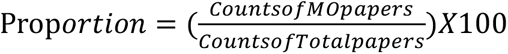

Temporal trends c assessed using linear regression and Pearson correlation analysis. For each MO–journal pair, we computed the Pearson correlation coefficient (r) between publication year and the corresponding proportion. Statistical significance of each correlation was evaluated via a two-tailed t-test, with significance levels indicated as: *, p<0.05;**, p<0.01; ***, p<0.001. Analyses were performed over four time windows: 1995–2024, and 1995–2004, 2005–2014, 2015–2024 for decade-specific trend evaluation. All analyses were performed in R (version 4.3.0). Data processing utilized the *tidyverse*, correlation and regression analyses were conducted using base R functions (*cor.test, lm*), and visualizations were generated with *ggplot2* and *pheatmap*. Code and derived datasets are available from the figshare repository (https://figshare.com/s/e79fdb15bbf1bd2074c4).

### Analysis of Conference Attendance Trends

Attendance records were obtained from major model organism conferences spanning the past several decades, including the International Worm Meeting (2011–2025), the Yeast Genetics Meeting (2012–2022), the Annual *Drosophila* Research Conference (2014–2025), the European *Drosophila* Research Conference (2013–2025) and the European Zebrafish Meeting (1999–2017). Data were provided by the respective organizing societies (the Genetics Society of America, the European *Drosophila* Society, and the European Zebrafish Society). Raw participant numbers were divided into six standardized role categories (except for the zebrafish meeting and the European *Drosophila* Research Conference, for which such role-wise breakdown was unavailable): Undergraduate, Graduate Student, Postdoc, Faculty/Lab Tech, Exhibitors/Staff/Speakers/Comps, and Others. Because these conferences are not held annually, years without available attendance data were excluded from analysis.

To assess temporal trends, we performed simple linear regression for each conference, modeling total attendance as a function of year. Additionally, separate linear regressions were conducted for each participant role to examine changes in its proportional representation over time. The strength and direction of each trend were quantified using the Pearson correlation coefficient (r) between calendar year and participant count (total or role-specific). Statistical significance of the correlations was evaluated with a two-tailed t-test, with p< 0.05 considered statistically significant. To assess whether changes in research output reflect shifts in community size or productivity per researcher, we compared publication trends with conference attendance data for each model organism. Conference attendance serves as a proxy for community size, under the assumption that stable or growing attendance alongside declining publications would indicate reduced productivity rather than a shrinking community. For each model organism, we generated dual-axis time series plots to visualize the relationship between annual publication counts and conference attendance over time. Years without available data for either metric were excluded from the analysis. We then quantified the linear relationship between attendance and publication count using Pearson correlation coefficients (r), with statistical significance evaluated as described above. A positive correlation would suggest that publication trends track community size, whereas a weak or negative correlation would point to changes in per-researcher productivity. Code and derived datasets are available from the figshare repository (https://figshare.com/s/e79fdb15bbf1bd2074c4).

### Analysis of shifts in publication preparation times

We analyzed changes in the time required for a scientific publication across years using data extracted from the NIH Reporter database. For each grant application ID, we used the fiscal year of an NIH grant award as the project start date, and the first paper published after the grant was awarded as the required time for a scientific publication. To extract the data, we manually downloaded tables for all *Drosophila* and Mouse grant applications from the NIH Reporter, then used the Python package pynih to access the NIH API and link each grant application ID to all published papers’ PMIDs associated with that grant. Then, using Biopython, we obtained information about each paper, such as the publication year. We then estimated the difference between each project start date and publication date. For papers with a negative span (sometimes papers that serve as a base for the grant get reported), we used the publication year closest to the project start date. We plotted the distribution per year of these differences to visualize a possible shift in the last 25 years. To quantify distribution shifts over time, we measured the mean for each year.

### Statistical analysis of publication trends by topics

Abstracts were classified into the categories “basic”, “applied”, “mechanistic” or “disease”, using the MoritzLaurer/deberta-v3-large-zeroshot-v2.0 model from (https://huggingface.co/MoritzLaurer/deberta-v3-large-zeroshot-v2.0). Our dataset contained 342,611 papers. 4668 papers did not have abstracts and were removed from the dataset. Each paper was passed through the model, which was set up to complete the hypothesis template “the scientific abstract refers to (2)” and given the classifiers “basic research”, “applied research”, “disease research” and “mechanisms of biological processes”. The terms “disease research” and “mechanisms of biological processes” were chosen as alternative indicators of applied and basic research since it wasn’t always clear if an abstract was applied or basic. The papers were simply categorized according to which was most likely, without a threshold for confidence. The average confidence score across all years was 69% and a range between 68% and 72% for each year. To test the quality of the classification we passed abstracts from the journals *Advances in Therapy*, which almost exclusively covers clinical medicine, and *Development* which mostly covers basic and mechanistic subjects (Fig. S16A, Table S3). We first ran the LLM on a collection of abstracts from these journals and then manually classified them to see how well the LLM lined up with a human judge. Taking all articles published in *Advances in Therapy* between 2000 and 2010 (N = 677), the model classified 605 (89%) as disease or applied research, 67 (10%) as basic or mechanistic and 5 (<1%) had no abstract. After human assessment of the abstracts classified basic or mechanistic, 56 (8%) were found to be misclassified disease or applied papers and 10 (1.5%) were correctly identified as basic or mechanistic. 1 abstract did not clearly state what the paper was about. Of the 605 papers classified as disease or applied, 10 (1.5%) were judged to be primarily basic or mechanistic. Conversely, to test its ability to identify basic and mechanistic papers, we passed abstracts from the journal Development, filtering for the keyword *shh* (for *sonic-hedgehog*, a developmental signaling gene) (Fig. S16B, Table S3). Out of all articles published between 1995 and 2024 (N = 473), 19 (4%) were classified as disease or applied research, 454 (96%) were classified as basic or mechanistic. After human assessment, 17 articles classified by the LLM as basic or mechanistic were judged to be disease or applied, 1 article classified as disease or applied was judged to be basic or mechanistic, and 1 article classified basic or mechanistic had an unclassifiable single-word abstract.

Given the subjectivity of this process, the exact numbers may vary and some papers could justifiably be fit into multiple categories. However, the overall trends appear to accurately reflect each journal’s profile Generally, the LLM seems to falsely classify disease or applied papers as basic or mechanistic—a false BM rate—at 11%. It seems to misclassify basic or mechanistic papers as applied or disease—a false AD rate—at 2.4%. This suggests a bias towards basic and mechanistic papers and the true number of disease and applied papers in our analysis may be around 10% higher than what we show. Although these journals were chosen to have a high number of clear-cut cases of either category, the full dataset likely contains many more borderline cases. Nevertheless, the trends in changing research profiles are likely real. Code and derived datasets are available from the figshare repository (https://figshare.com/s/e79fdb15bbf1bd2074c4). Table S2 is available from the figshare repository (https://figshare.com/s/e959259f1f4bd22ad9be?file=63304267).

### Analysis of Funding Trends

ERC funding data were retrieved from the Dashboard of ERC funded projects and evaluated proposals (https://erc.europa.eu/projects-statistics/erc-dashboard). Only three ERC grant schemes were retained for analysis: Starting Grants, Consolidator Grants, and Advanced Grants. NIH funding data were retrieved from NIH RePORTER (https://reporter.nih.gov/); all NIH award types were included in the analysis. Both databases support keyword searches across the title and abstract of funded projects, and for NIH RePORTER ‘scientific terms’. We used the same search key words for MOs and emerging research areas (including AI, cross-disciplinary studies, and organoids) that were applied in our literature search, and quantified the funding allocated to matching projects both in absolute amount and as a share of total research funding.

## Acknowledgements

We are grateful to several colleagues who provided constructive feedback at different stages of this project, also helping us identify literature and data, namely: Richard Benton, Nick Brown, Virginie Courtier, Bart Deplancke, Eileen Furlong, Mark Hanson, Julie Hayner Simpson, Nikolaos Konstantinides, Artyom Kopp, Peter Lawrence, Bruno Lemaitre, Irene Miguel Aliaga, Marco Milan, Kate O’Connor-Giles, Luisa F. Pallares, Thomas Préat, Andreas Prokop, Todd Schlenke, Gilles Storelli, Miltos Tsiantis, Trisha Wittkopp, Helena Cocheme, Lukas Neukomm, Benjamin Prud’homme. We are particularly grateful to Suzy Brown (Genetics Society of America), Stefan Schulte-Merker (the European Zebrafish Society), Bruno Lemaitre, Javier Morante Oria, François Leulier and Nic Tapon (European *Drosophila* Society) for sharing and letting us use meeting attendance information.

---

**Figure S1.**
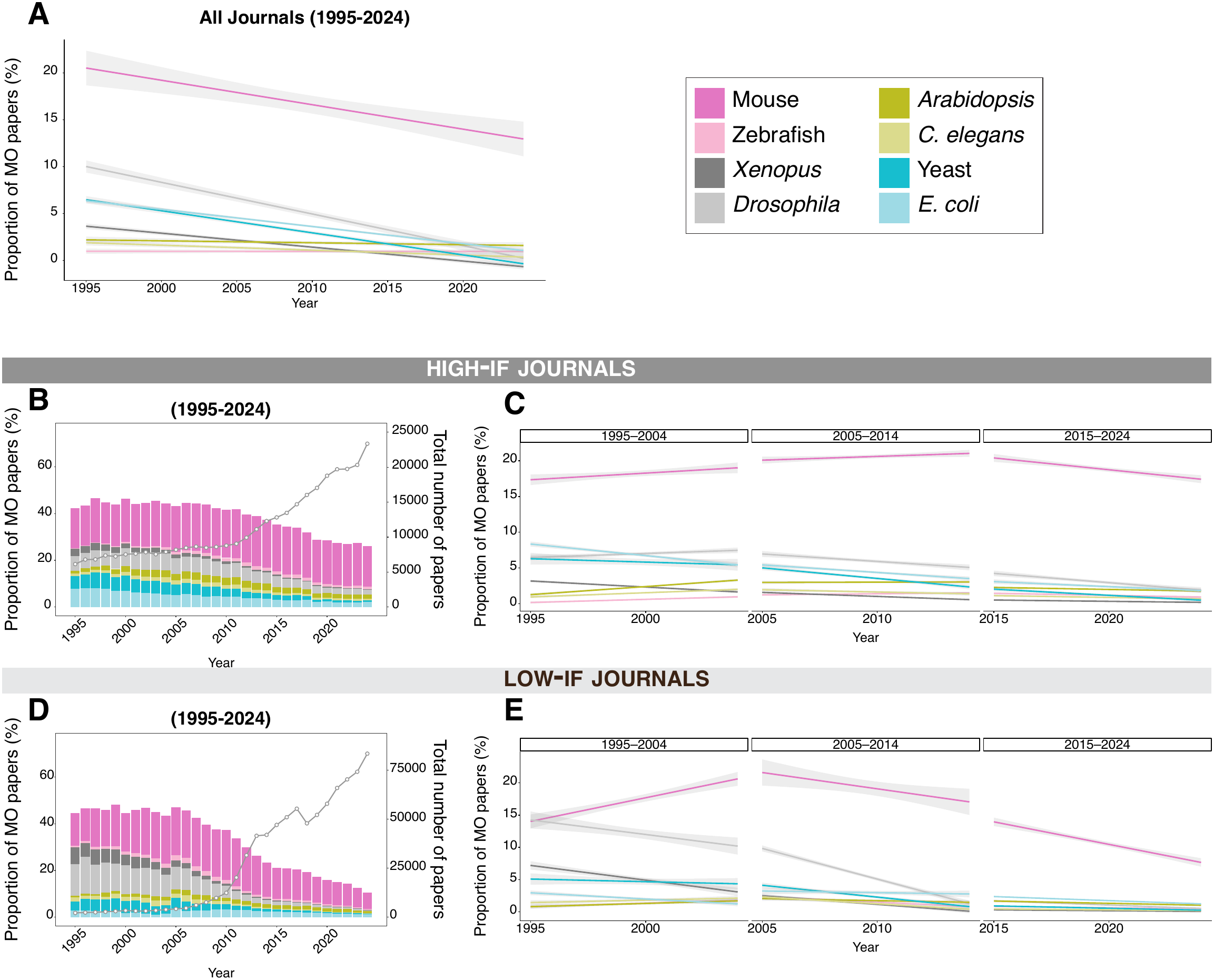
Representation of MO research over time by journal impact factor. **A**. Linear regression lines with confidence intervals (shaded areas) indicate publication trends for selected journals listed in Table S1 for the period 1995–2024. Each line represents one model organism. Pearson correlation coefficients (r) and their statistical significance (*) are as follows: mouse: r = -0.68***; zebrafish: r = -0.02; *Xenopus*: r = -0.96***; *Drosophila*: r = -0.96***; *Arabidopsis*: r = -0.36; *C. elegans*: r = -0.76***; yeast: r = -0.97***; *E. coli*: r = -0.98***. **B**. Proportion (%) of papers using different model organisms across high-IF journals from 1995 to 2024. Each colored bar or trend line represents a specific model organism. **C**. Linear regression lines with 95% confidence intervals indicate publication trends in high-IF journals over time. **D**. Proportion (%) of papers using different model organisms across low-IF journals from 1995 to 2024. **E**. Linear regression lines with 95% confidence intervals indicate publication trends in low-IF journals over time. Corresponding Pearson correlation coefficients and statistical significance are summarized in Tables S5–S7.

**Figure S2.**
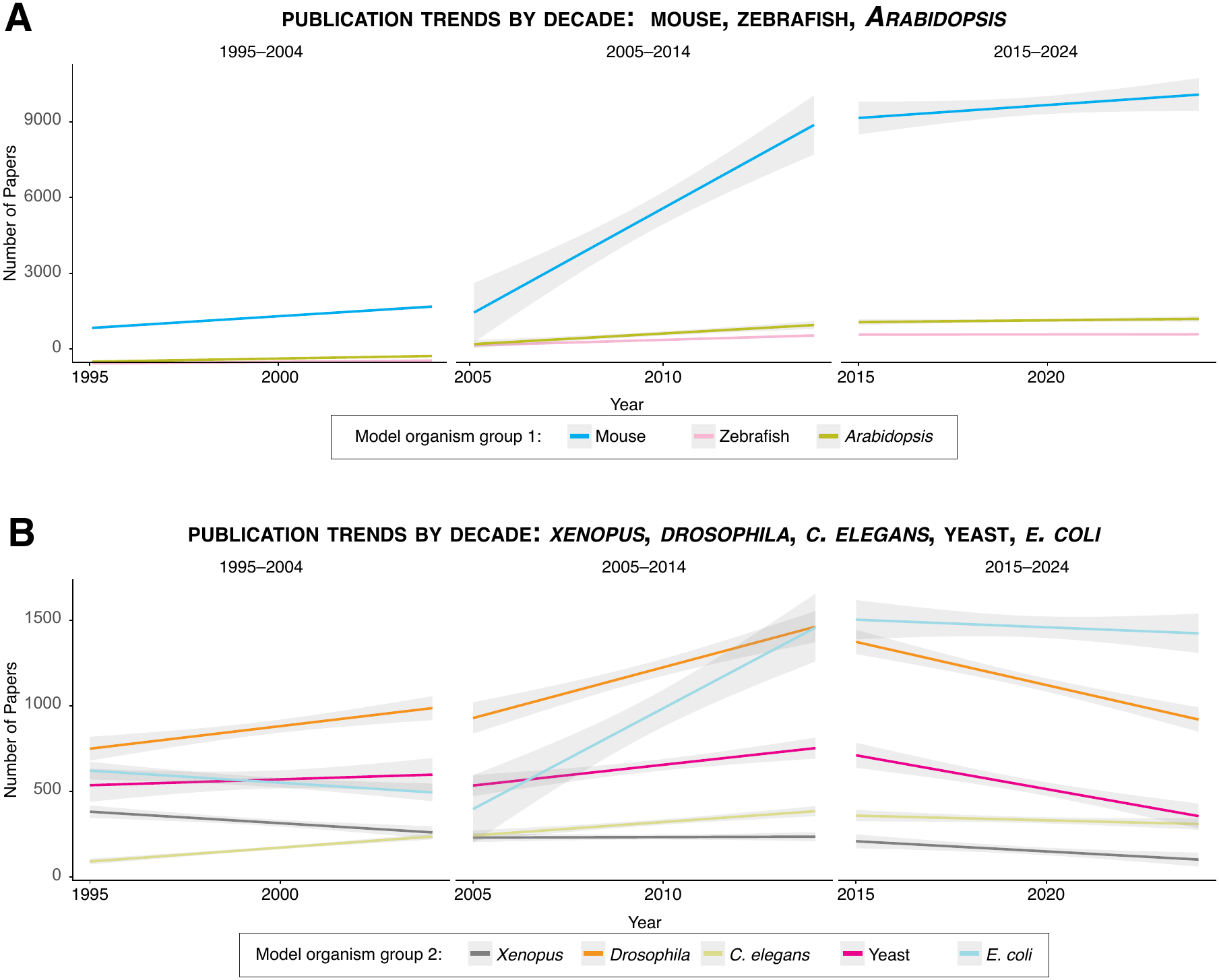
Decade-wise trends in publication counts across all journals. Linear regression lines with confidence intervals (shaded areas) indicate publication trends over time for two groups of model organisms: mouse, zebrafish and *Arabidopsis* (**A**) and *Xenopus*, *Drosophila*, *C. elegans*, yeast and *E. coli* (**B**). Each line represents one model organism. Pearson correlation coefficients (r) and their statistical significance (*) are as follows: mouse: r = -0.68***; zebrafish: r = -0.02; *Xenopus*: r = -0.96***; *Drosophila*: r = -0.96***; *Arabidopsis*: r = -0.36; *C. elegans*: r = -0.76***; yeast: r = -0.97***; *E. coli*: r = -0.98***. Linear regression lines with 95% confidence intervals show decade-specific publication trends for each model organism (1995–2004, 2005–2014, 2015–2024). Corresponding Pearson correlation coefficients and statistical significance are summarized in Table S9.

**Figure S3.**
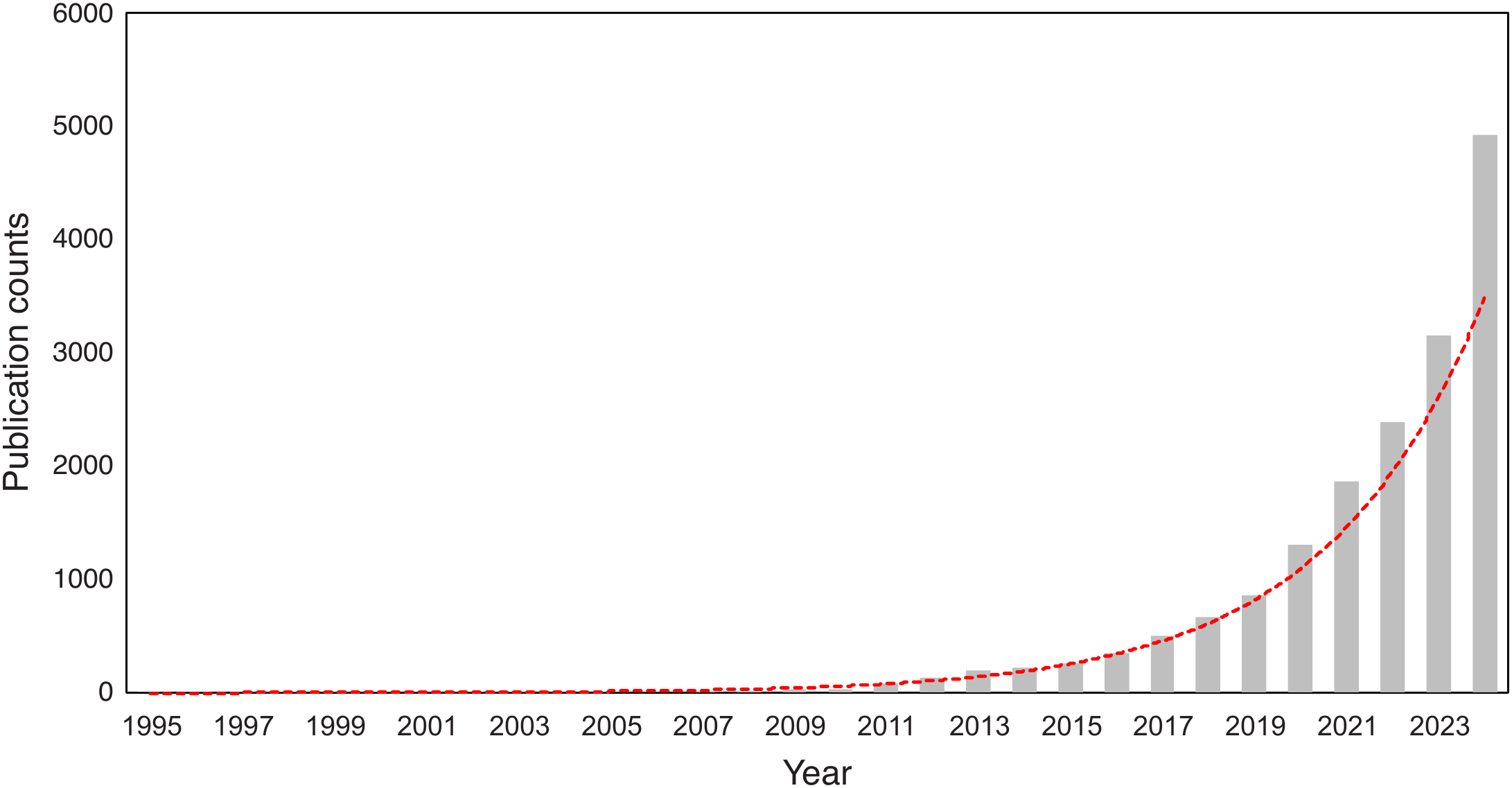
Trend in publication counts for emerging research topics across selected journals, 1995–2024. Publications were identified by searching the Web of Science for articles related to cross-disciplinary research, non-model organisms, organoids, AI, or machine learning, after excluding all model organism-focused articles. The red line illustrates the exponential growth in the annual number of publications on these topics over the period. Fit curve (red): y = 0.5971 e^0.2891x^; R^2^ = 0.9918.

**Figure S4.**
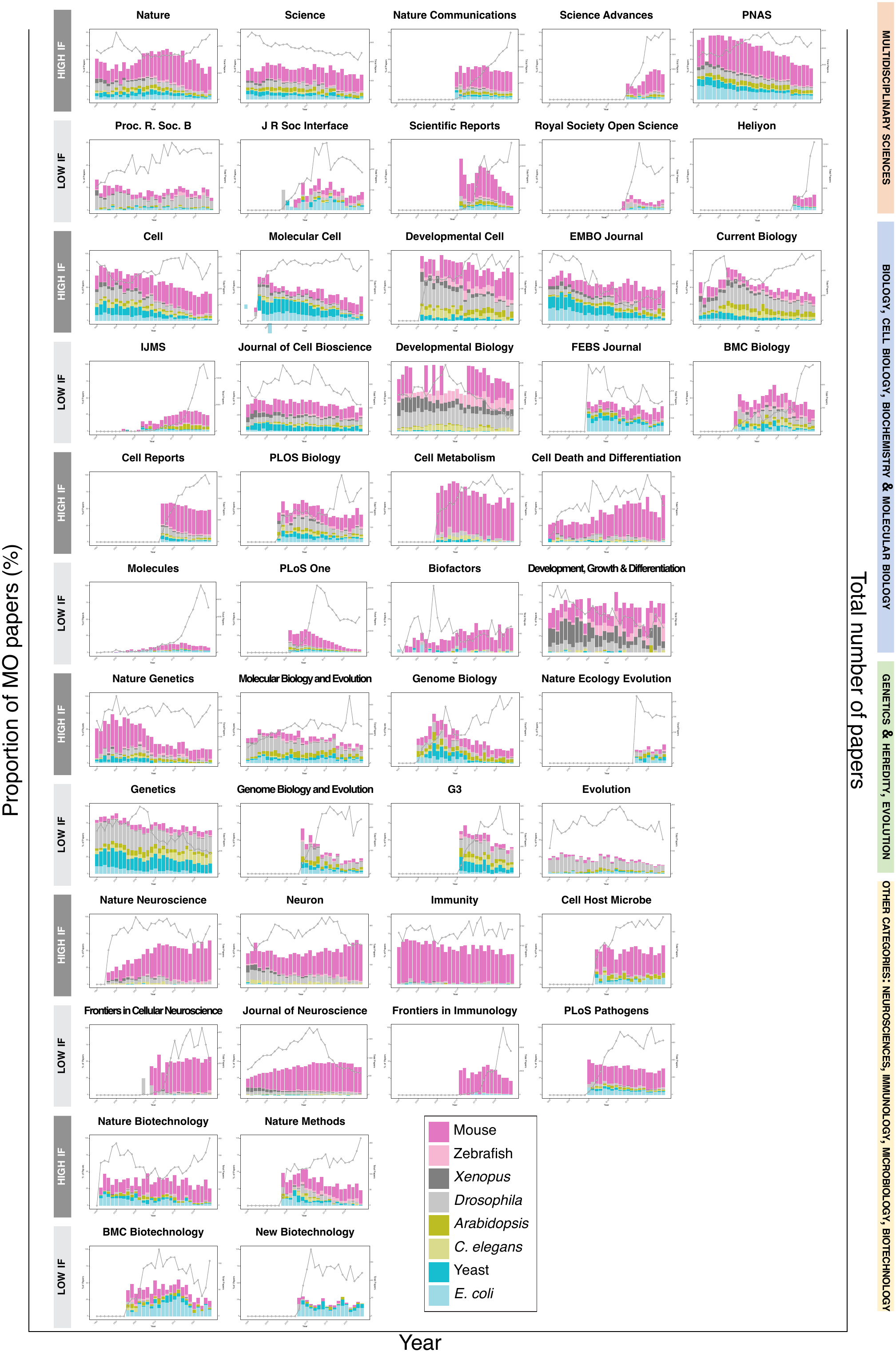
Publication trends of different model organisms in 48 individual journals. Proportion (%) of papers using different model organisms in 48 selected journals from 1995 to 2024. Each high-IF journal (upper plot) can be compared to a low-IF counterpart (lower plot). Each colored bar represents a specific model organism.

**Figure S5.**
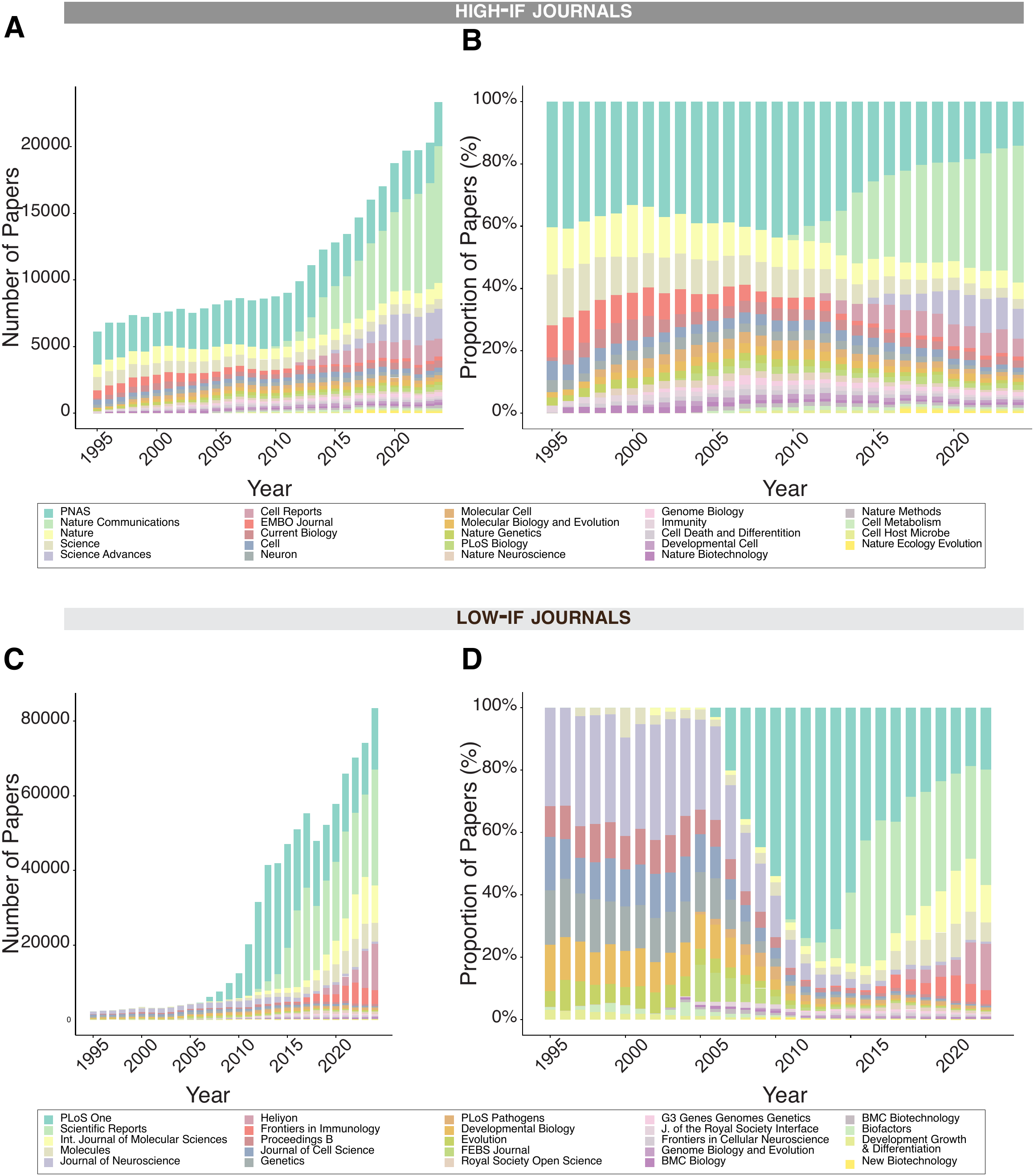
Publication output of selected journals over time. **A–B.** High-impact-factor journals. Stacked bar plots show (**A**) Annual publication counts of selected high-IF journals from 1995 to 2024. (**B**) Proportional contribution (%) of each journal to the total publications within this category over the same period. Each colored segment representing one journal. **C–D.** Low-impact-factor journals. Stacked bar plots show (**C**) Annual publication counts of selected low-IF journals. (**D**) Proportional contribution (%) of each journal within the low-IF category. Each colored segment representing one journal.

**Figure S6.**
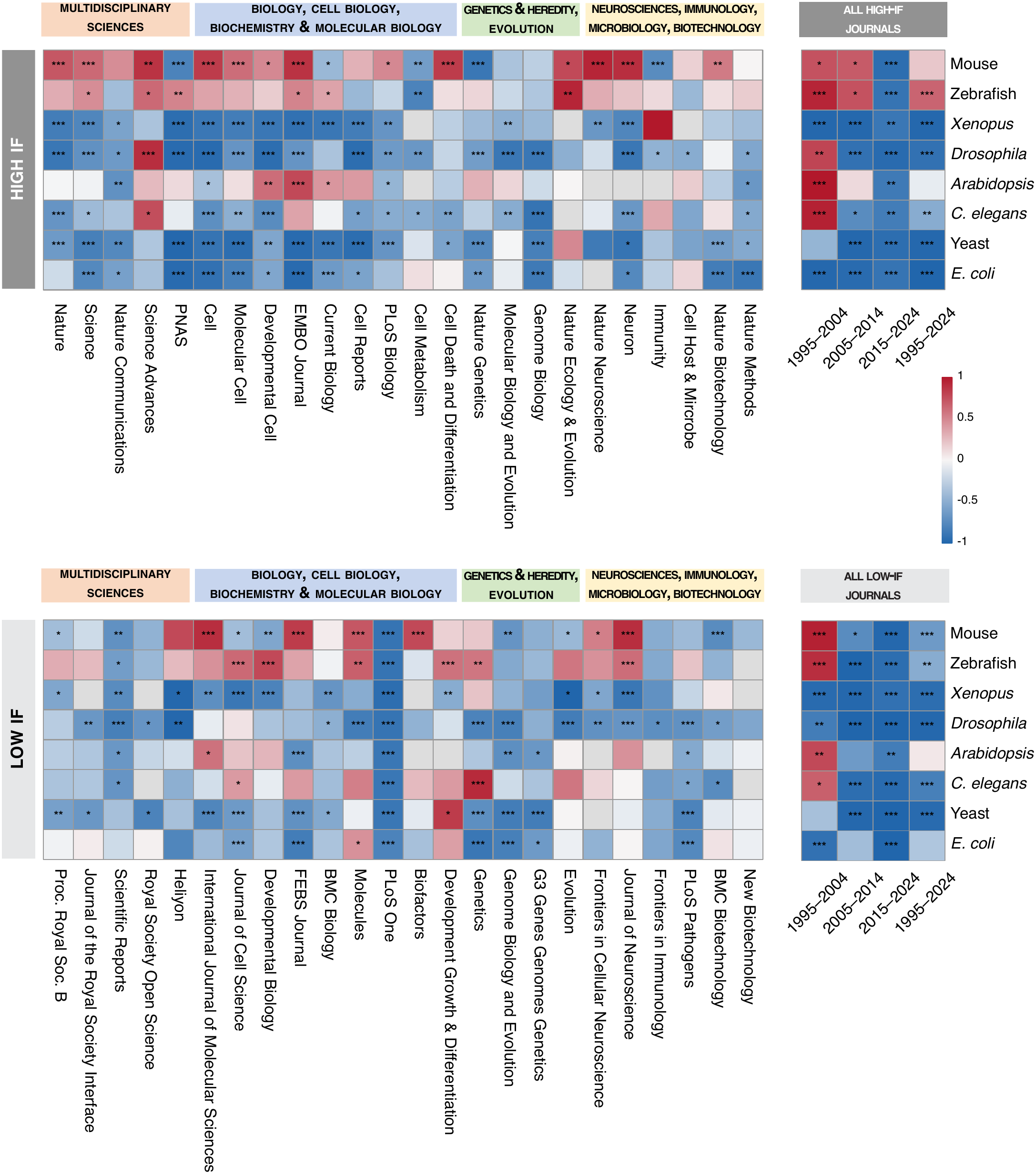
Correlation heatmap of publication trends for model organisms in all selected journals. Pearson correlation coefficients (r) between publication year (Left panel: year 1995 to 2024; Right panel: 4 time periods comparison) and proportion of papers was calculated for eight model organisms across selected journals. Each cell represents the correlation strength for a specific organism-journal combination, with blue indicating decreasing trends (r < 0), white indicating no correlation, and red indicating increasing trends (r > 0).

**Figure S7.**
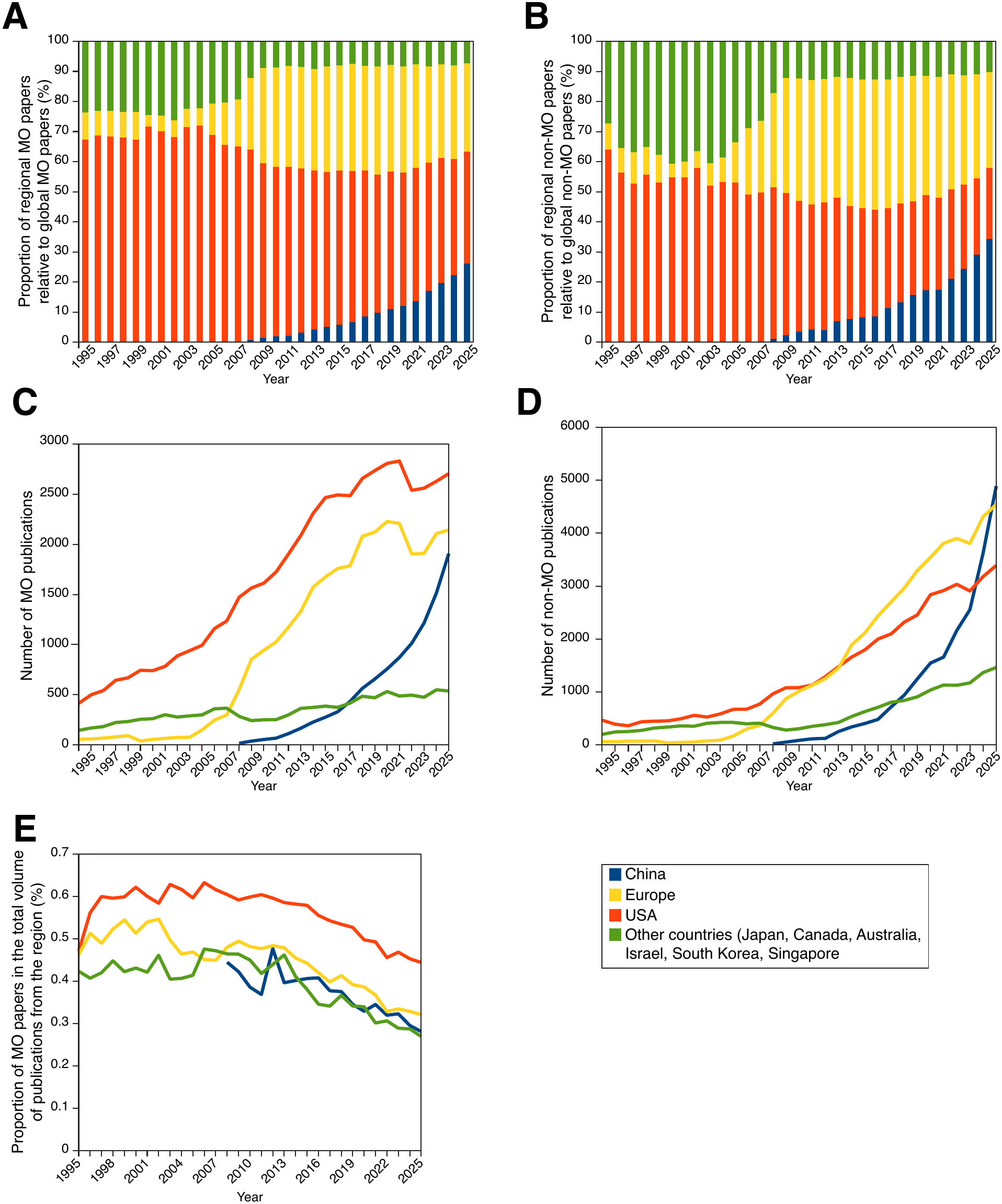
Regional trends in MO and non-MO publications across 24 high-impact journals over the past three decades. **A–B**. Changes in the proportional representation of MO (**A**) and non-MO (**B**) papers across four regions: China, the USA, Europe, and Others. Each region’s share is calculated relative to the total number of papers in that category worldwide. **C–D**. Changes in the annual publication volume of MO (**C**) and non-MO (**D**) papers by region over the same period. (**E**) Region-specific trends in the proportion of MO papers within each region’s own total output, calculated as the number of MO papers published by a given region in a given year divided by the total number of papers published by that same region in that year. This panel reflects how the relative weight of MO research has changed within each region over time.

**Figure S8.**
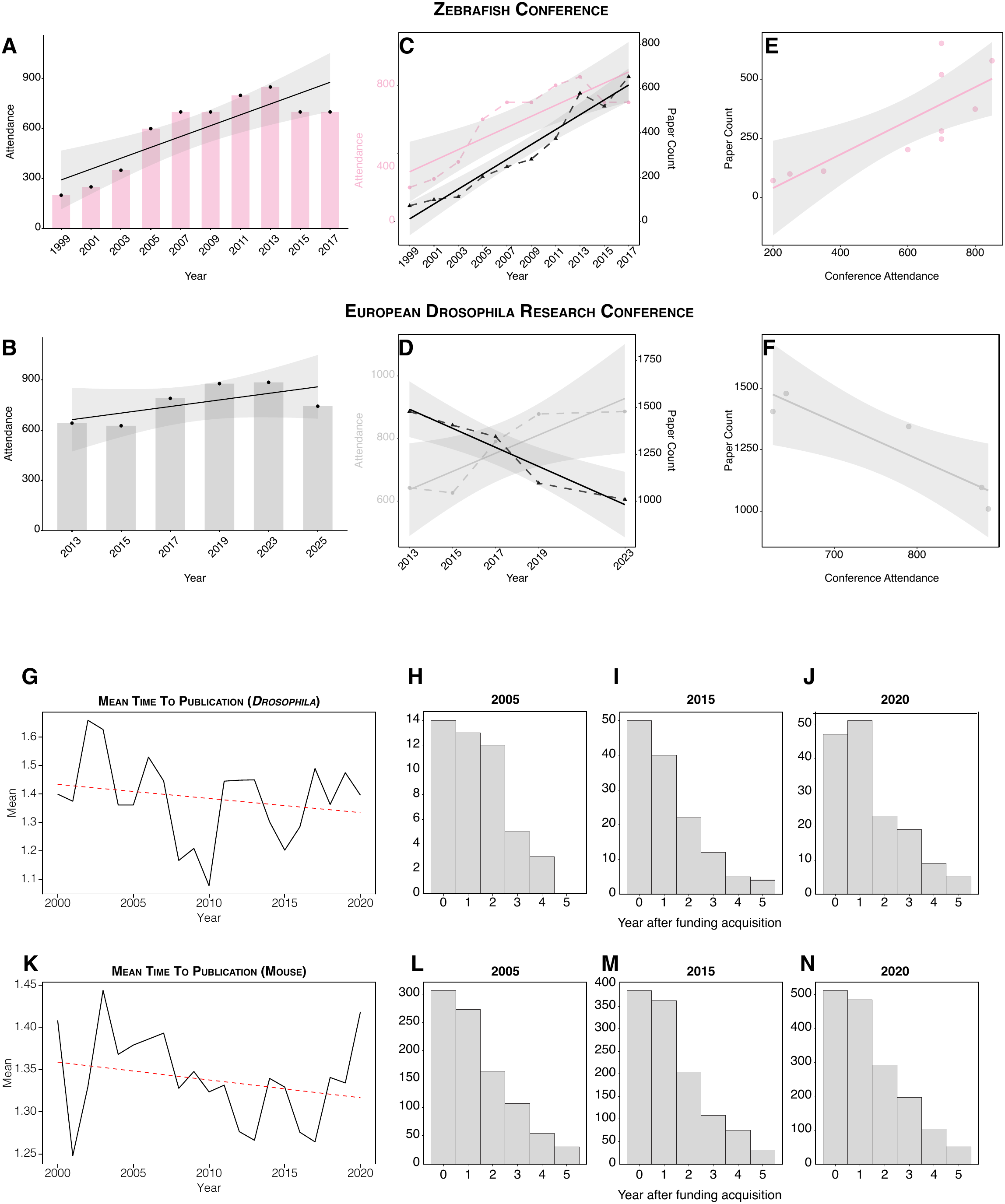
Conference attendance and temporal publication trends for MOs. **A–B.** Number of participants at European conferences for zebrafish (A) and *Drosophila* (B). **C–D.** Dual-axis time series plots showing annual changes in conference attendance (colored lines, left y-axis) and publication output (black lines, right y-axis) for *Drosophila* (**C**) and zebrafish (**D**). Solid lines represent linear trends, with shaded areas indicating 95% confidence intervals. **E–F.** Correlation between annual conference attendance and publication count for each model organism. Pearson correlation coefficients (r) and p-values are as follows: (**E**) zebrafish: r = 0.79, p = 0.007; (**F**) *Drosophila* (EDRC): r = –0.92, p = 0.02. **G–J.** Mean for delay between grant award and first resulting publication within a 5-year window for *Drosophila*. Temporal trends (**G**) (r=-0.21, p=0.35), and distributions using a 5-year window after a project’s start date for 2005 (**H**), 2015 (**I**) and 2020 (**J**). **K–N.** Mean for delay between grant award and first resulting publication within a 5-year window for mouse. Temporal trends (**K**) (r=-0.24, p=0.29), and distributions using a 5-year window after a project’s start date for 2005 (**L**), 2015 (**M**) and 2020 (**N**).

**Figure S9.**
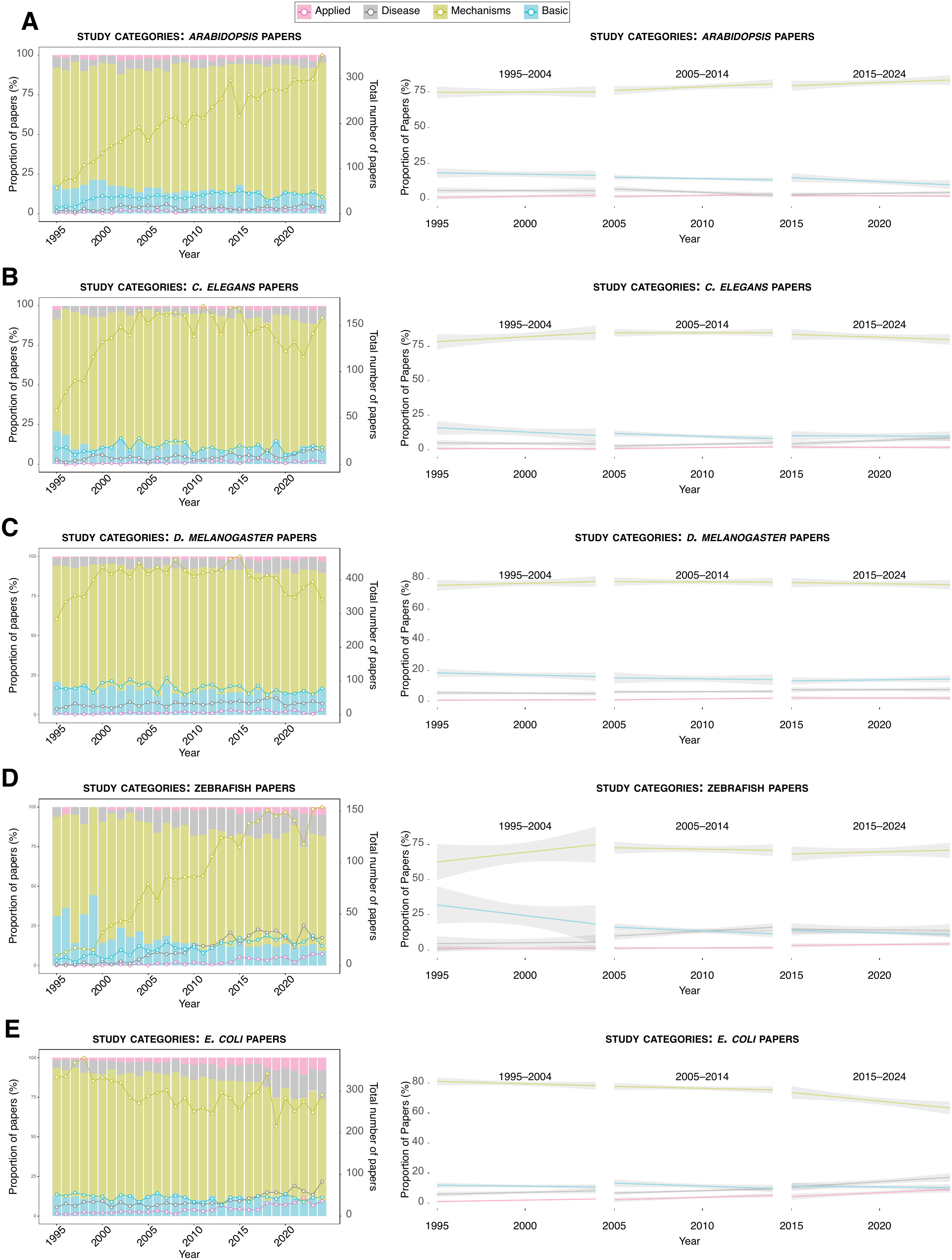
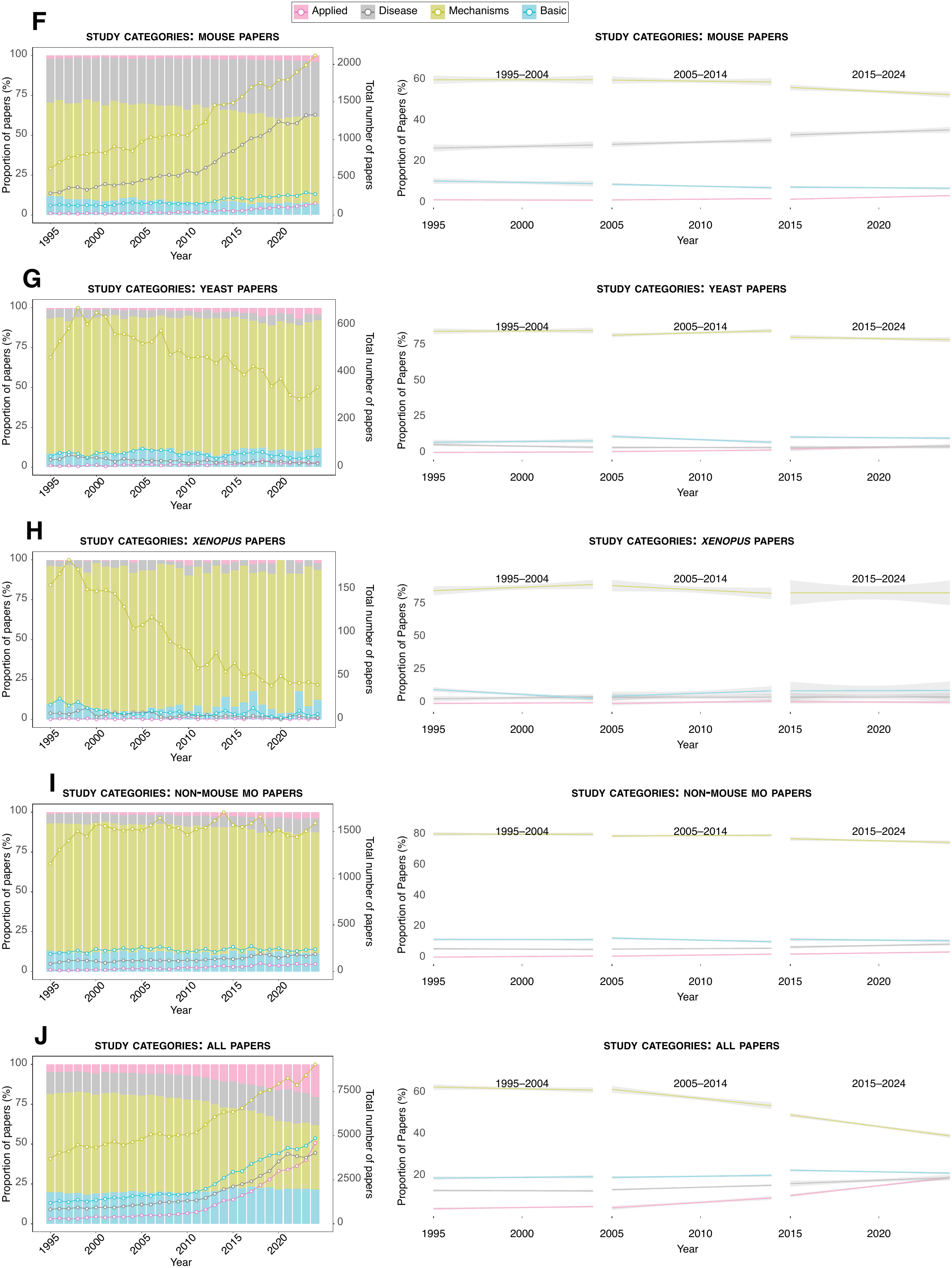
Type of research papers in high-IF journals from the categories ‘basic’, ‘mechanistic’, ‘applied’ and ‘disease’ according to an LLM classifier based on abstracts for every model organism individually. **A.** (left) *Arabidopsis* papers of each category published per year from 1995–2024 (left axis, proportion of papers for each year; right axis, number of papers). (right) Change in proportion of *Arabidopsis* papers of each category published per year over the decades 1995–2004, 2005–2014 and 2015–2024. B. Same for *C. elegans.* C. Same for *D. melanogaster.* D. Same for zebrafish. E. Same for *E. coli.* F. Same for mouse. G. Same for yeast. H. Same for *Xenopus.* I. Same for non-mouse model organisms. **J**. Same for all papers.

**Figure S10.**
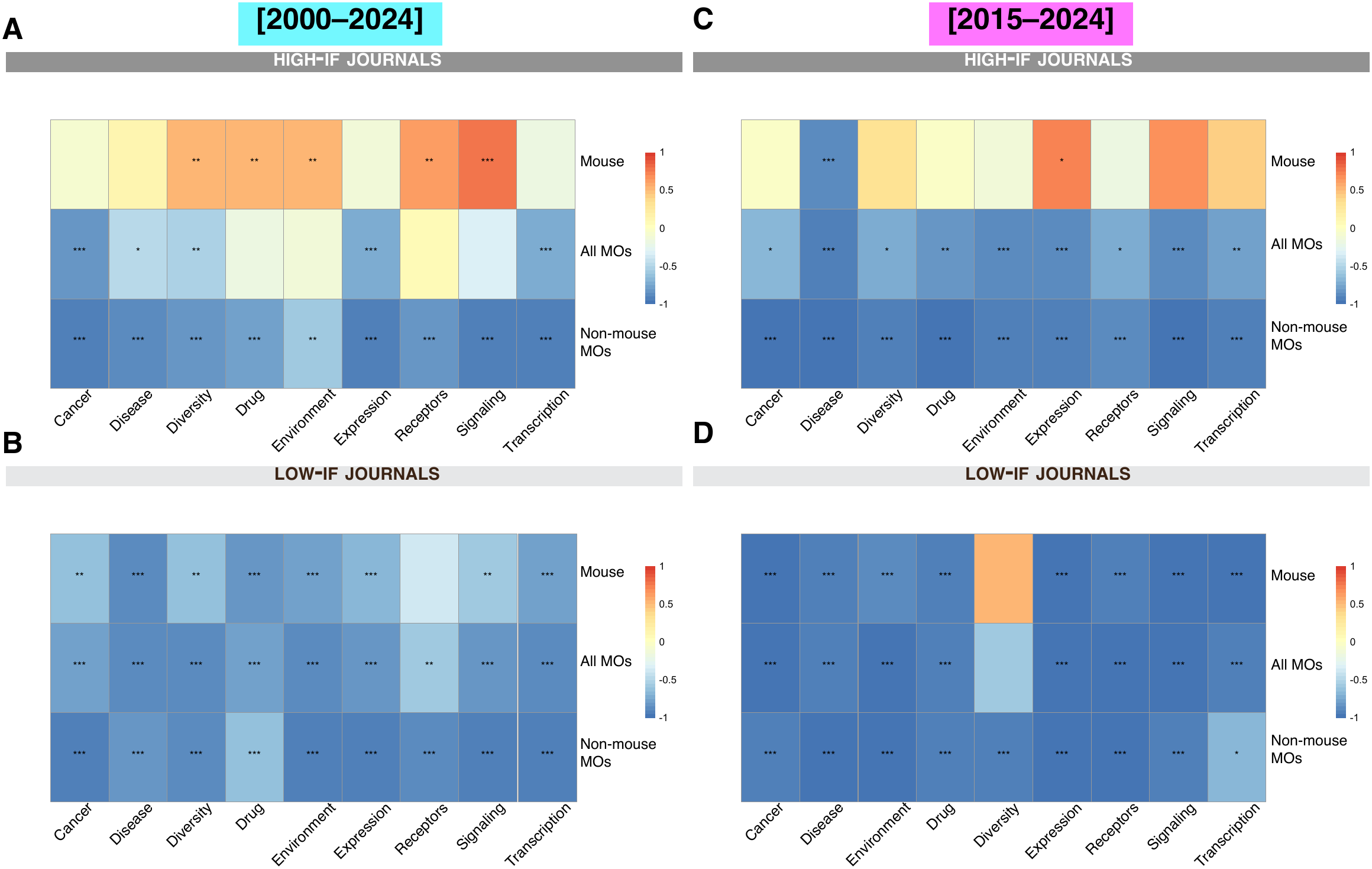
Specific keyword trends in high-IF and low-IF journals between over different time periods. **A,C**. High-IF journals. **B, D**. Low-IF journals. Correlation heatmaps for the evolution of the proportion of papers that include the selected keyword (cancer, disease, diversity, drug, environment, expression, receptors, signaling, or transcription) from 2000–2024 (**A–B**) or from 2015–2024 (**C–D**). Pearson correlation coefficients (r) were estimated for each of the specific keywords for papers including all MOs, mouse only, and non-mouse-MO research. Blue indicates decreasing trends (r < 0), white indicates no correlation, and red indicates increasing trends (r > 0), as shown on the colormap. Statistical significance is indicated by asterisks within each cell.

**Figure S11.**
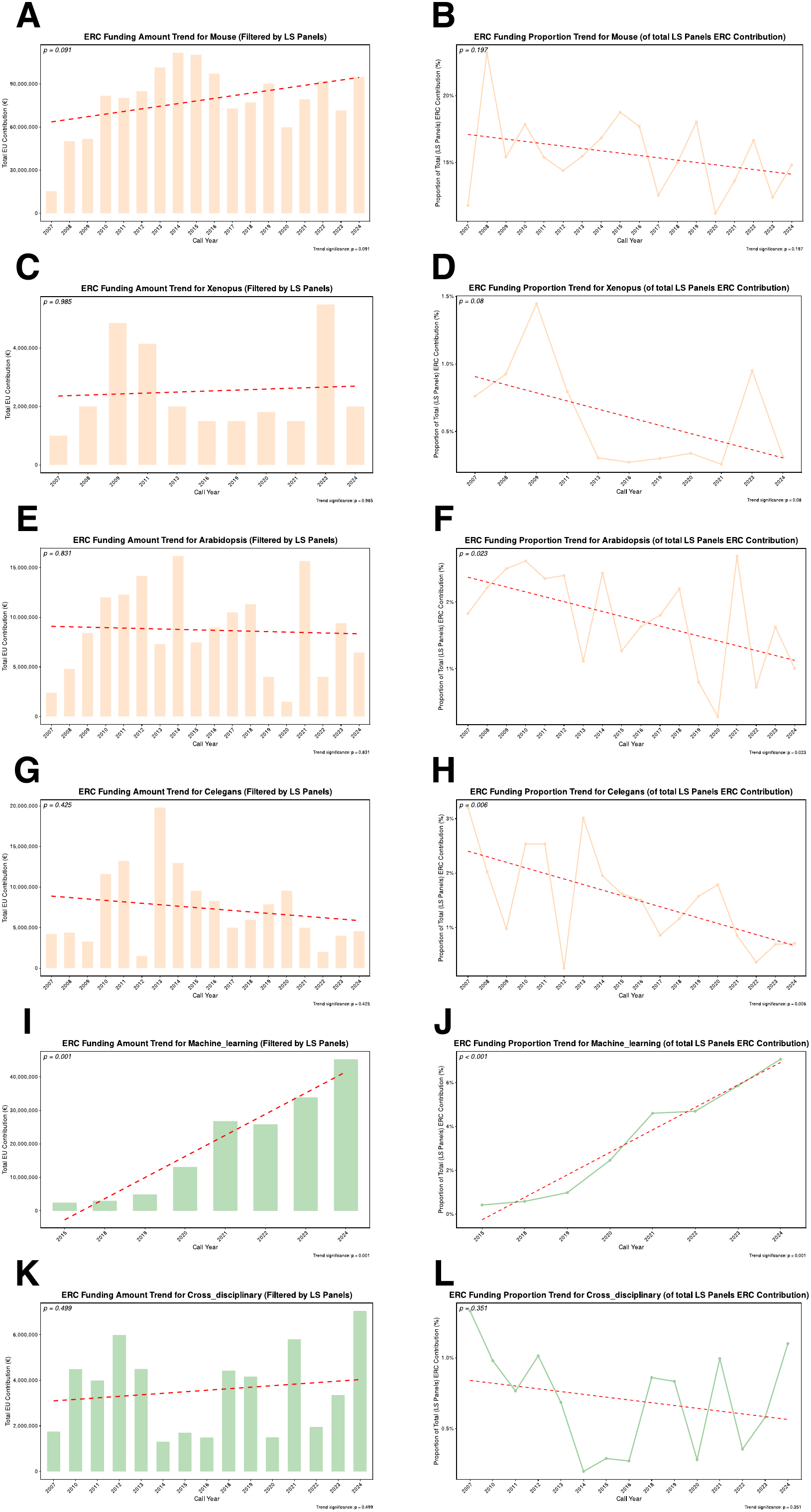
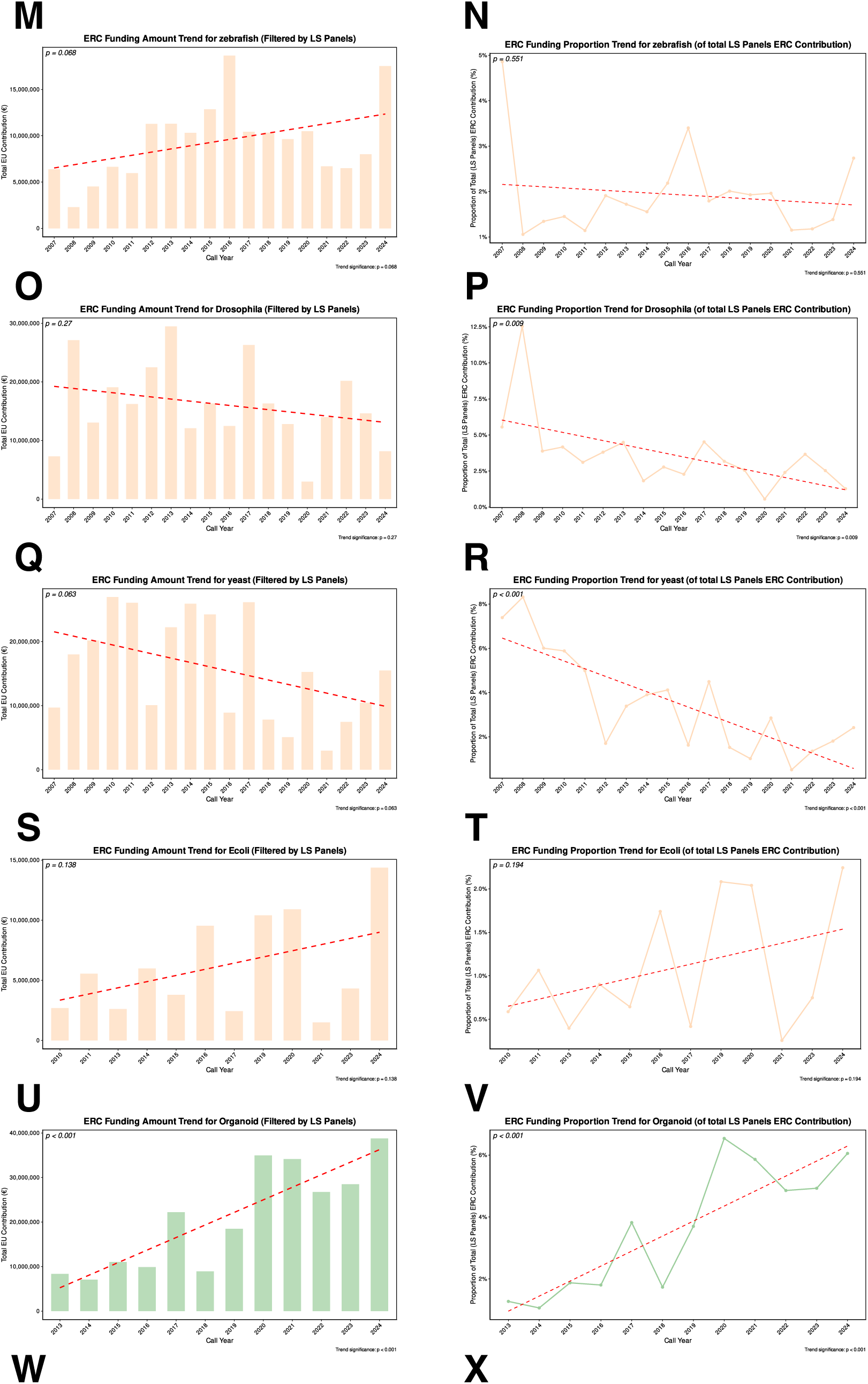
ERC funding (LS panels) trends for classical model organisms and emerging fields. For each category, the left panel shows the total amount of ERC funding for LS panels linked to a given topic (in M€), the right panel shows that amount as a proportion of total ERC research funding for LS panels per year (**A–B**) mouse, (**C–D**) *Xenopus* (**E–F**) *Arabidopsis* (**G–H**) *C. elegans*, (**I–J**) machine learning, (**K–L**) cross disciplinary, (**M–N**) zebrafish, (**O–P**) *Drosophila*, (**Q–R**) yeast, (**S–T**) *E. coli*, (**U–V**) organoid.

**Figure S12.**
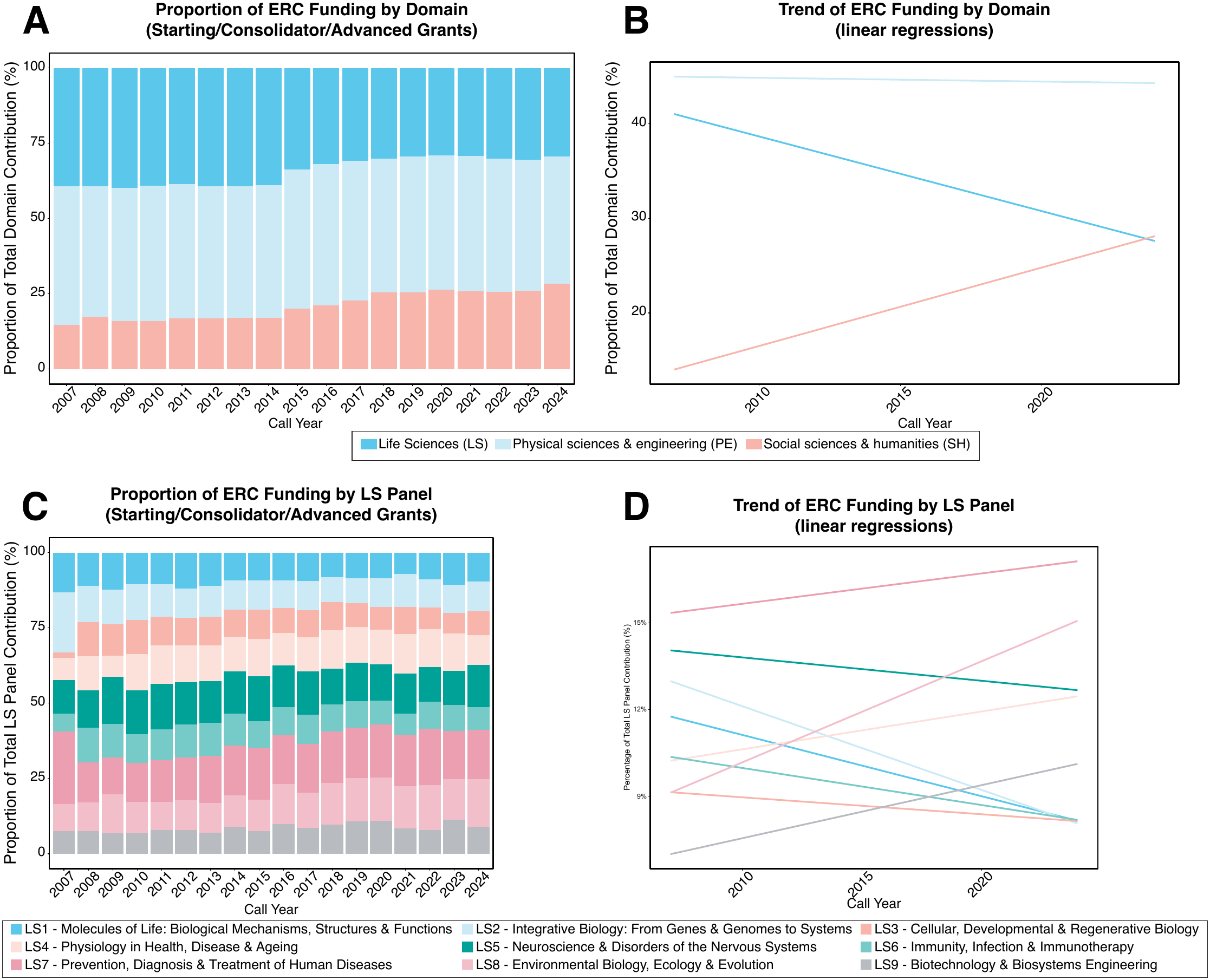
Temporal trends in ERC funding distribution across scientific domains. **A–B**. Changes in the relative distribution of ERC funding across the three main scientific domains: Life Sciences (LS), Physical Sciences & Engineering (PE), and Social Sciences & Humanities (SH). **A**. Stacked bar plot showing the proportional composition of funding across domains over time. **B**. Linear trends illustrating the change in funding proportion for each domain across the study period. **C–D**. Changes in the distribution of ERC funding within the Life Sciences domain, broken down by its eight sub-panels (LS1-LS8). **C**. Stacked bar plot showing the proportional composition of funding across LS sub-panels over time. **D**. Linear regression lines depicting the direction and magnitude of funding proportion changes for each LS sub-panel across the study period.

**Figure S13.**
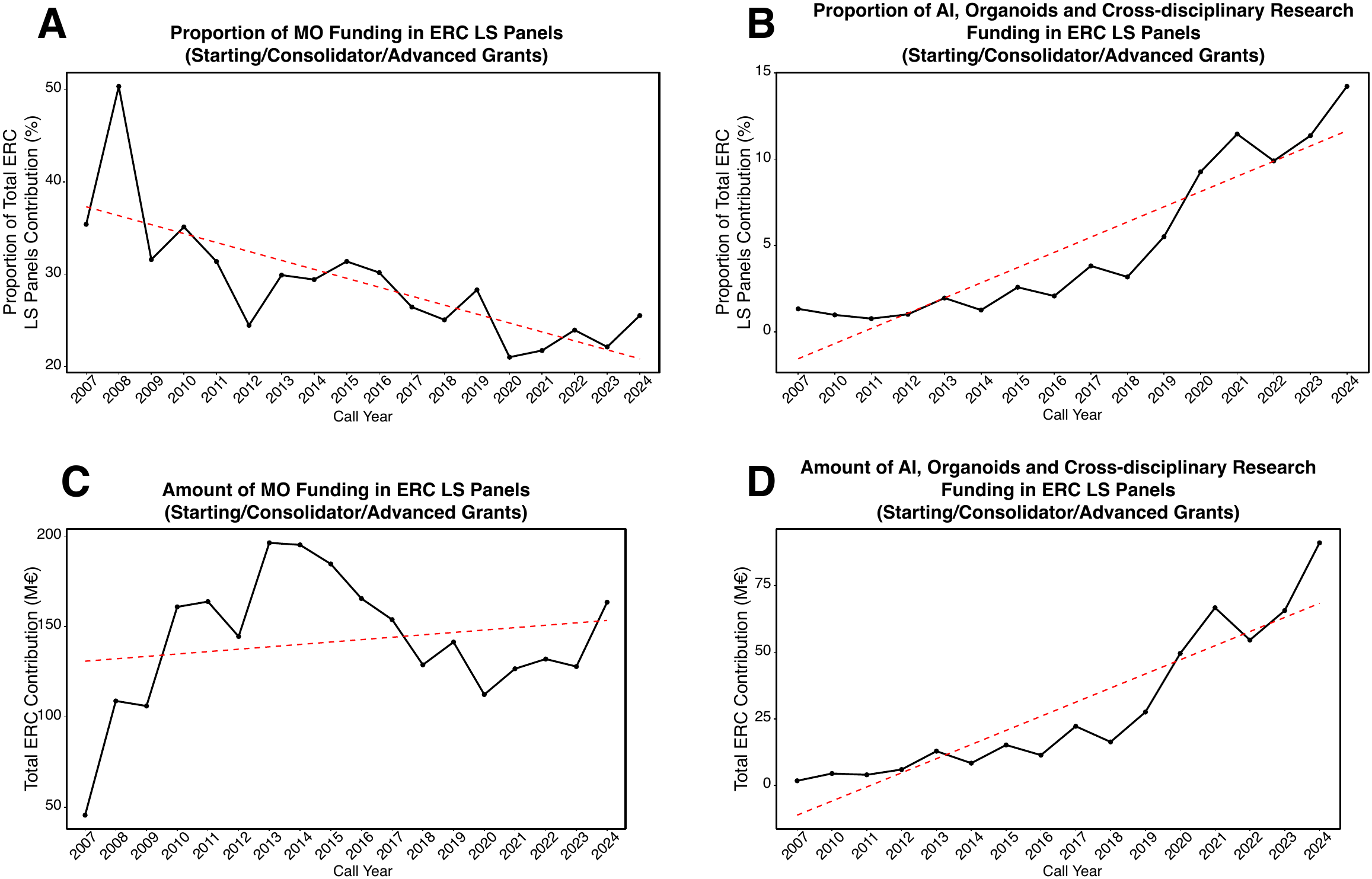
ERC funding trends for classical model organisms and emerging fields. **A–B.** Proportion of total ERC funding for classical model organisms (**A**) and emerging fields (**B**). **C–D**. Absolute funding amounts (M€) for classical model organisms (**C**) and emerging fields (**D**). Emerging fields include machine learning, organoids, and cross-disciplinary research.

**Figure S14.**
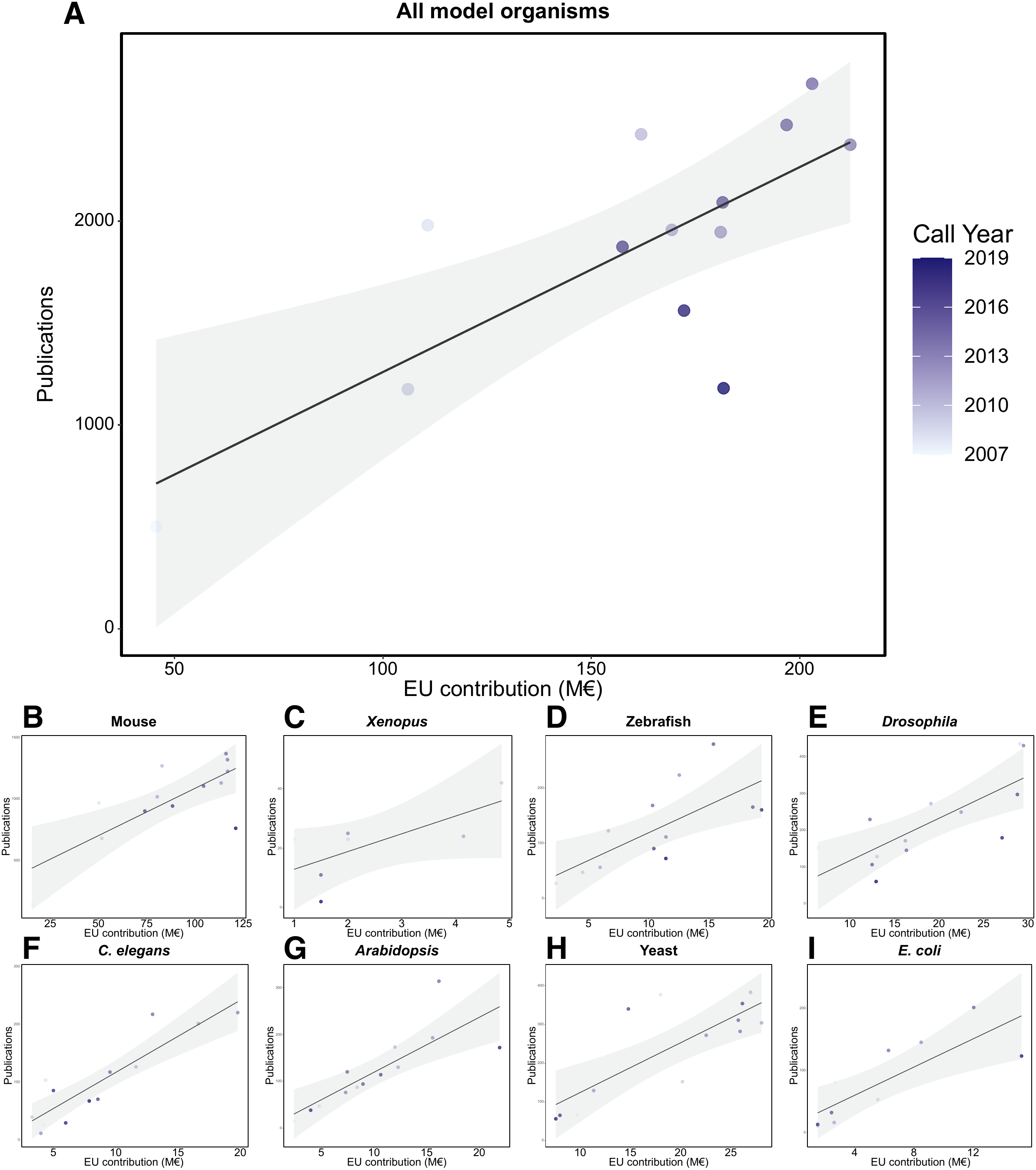
Association between ERC funding and publication output for classical model organism projects. **A**. All classical MO projects pooled: relationship between project funding (M€) and number of resulting publications. **B–I**. Per MO species: relationship between project funding (€) and publication count, shown separately for mouse (**B**), *Xenopus* (**C**), zebrafish (**D**), *Drosophila* (**E**), *C. elegans* (**F**), *Arabidopsis* (**G**), yeast (**H**) and *E. coli* (**I**).

**Figure S15.**
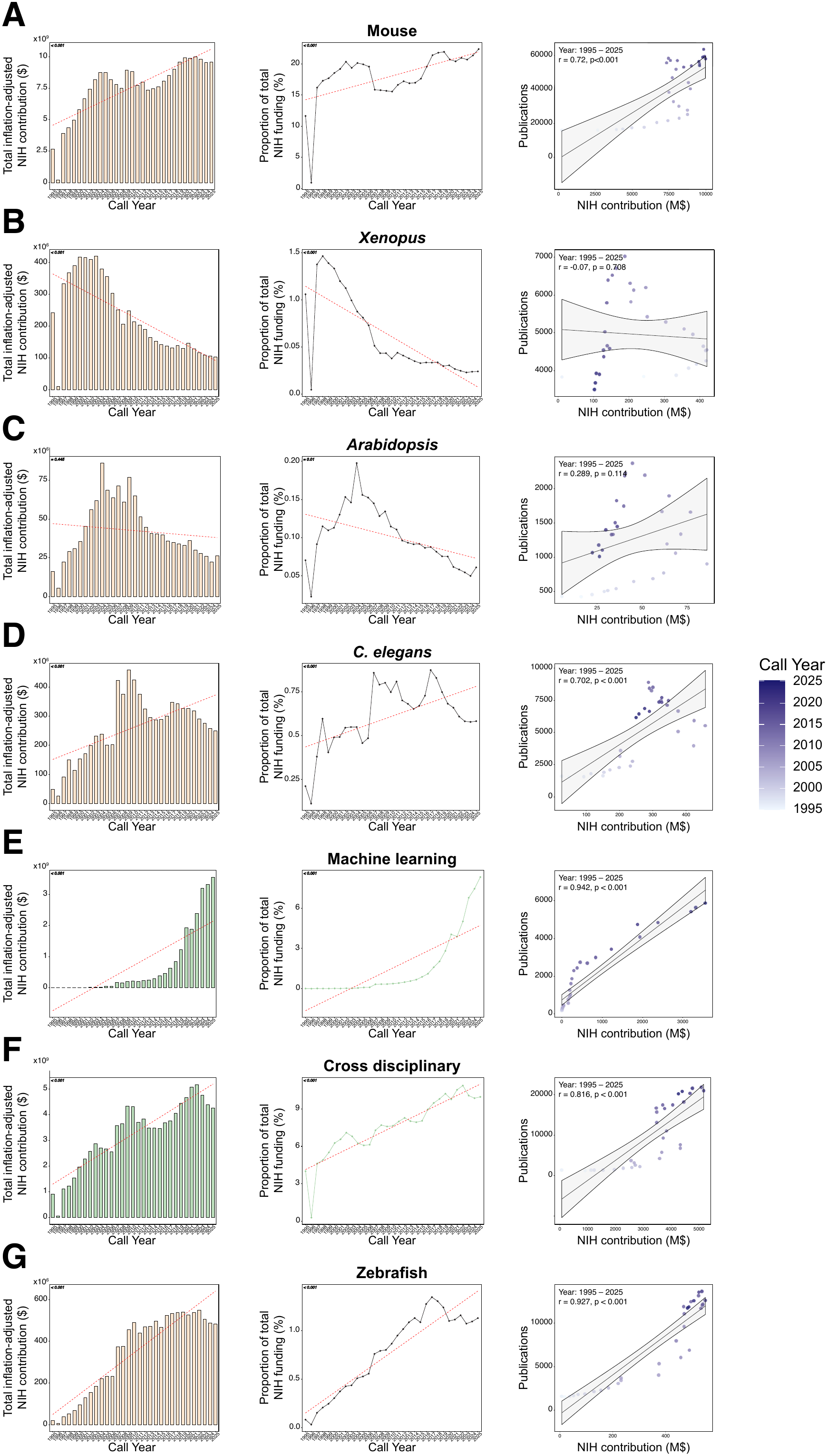
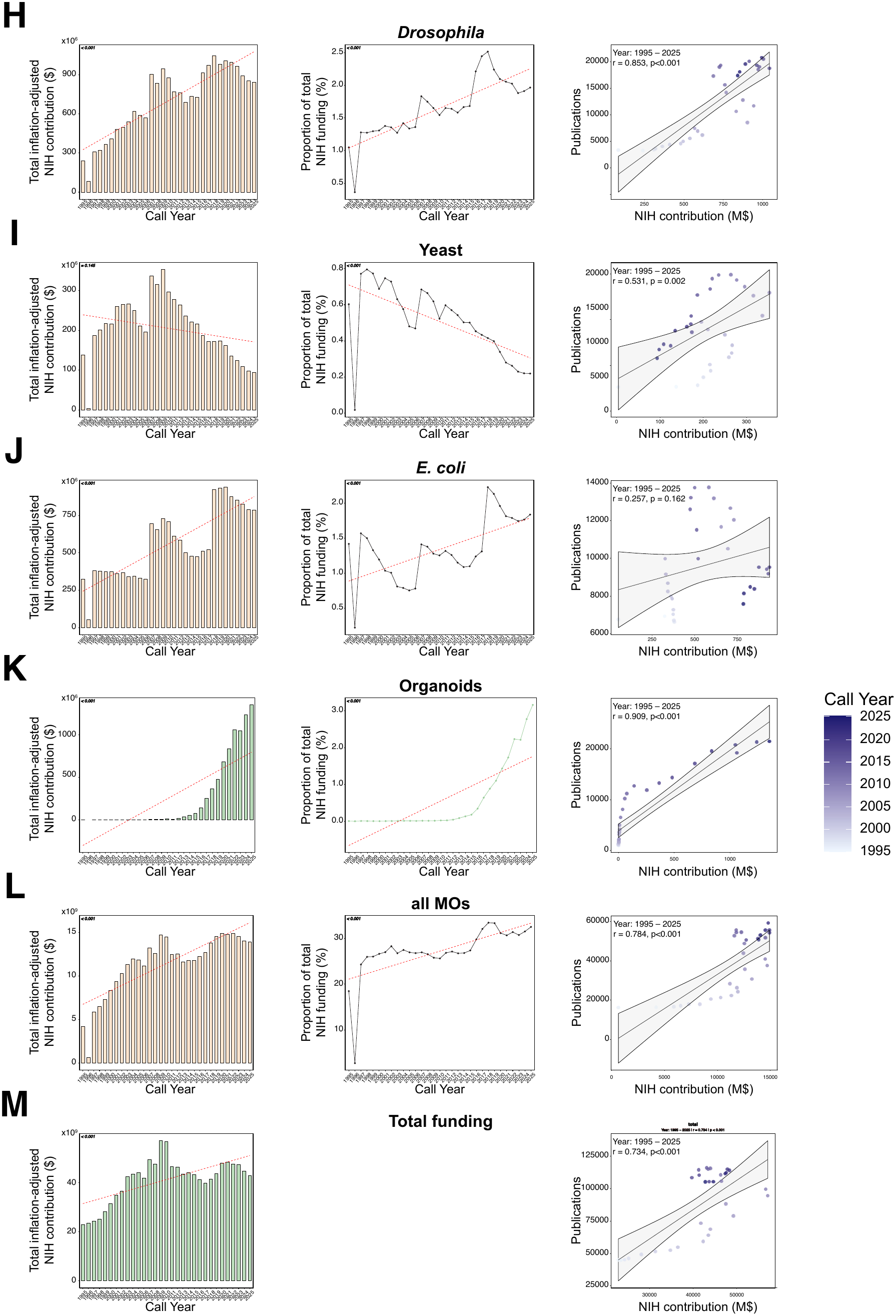
NIH funding trends for classical model organisms and emerging fields. For each category, the left panel shows the total amount of NIH funding linked to a given topic (in $ adjusted to 2025 levels using the biomedical research and development price index), the middle panel shows that amount as a proportion of total NIH research funding per year, and the right panel shows the correlation between funding and publications produced with NIH funding. mouse (**A**), *Xenopus* (**B**), *Arabidopsis* (**C**), *C. elegans* (**D**), machine learning (**E**), cross disciplinary (**F**), zebrafish (**G**), *Drosophila* (**H**), yeast (**I**), *E. coli* (**J**), organoid (**K**), All MOs (**L**), total NIH research funding (**M**).

**Figure S16.**
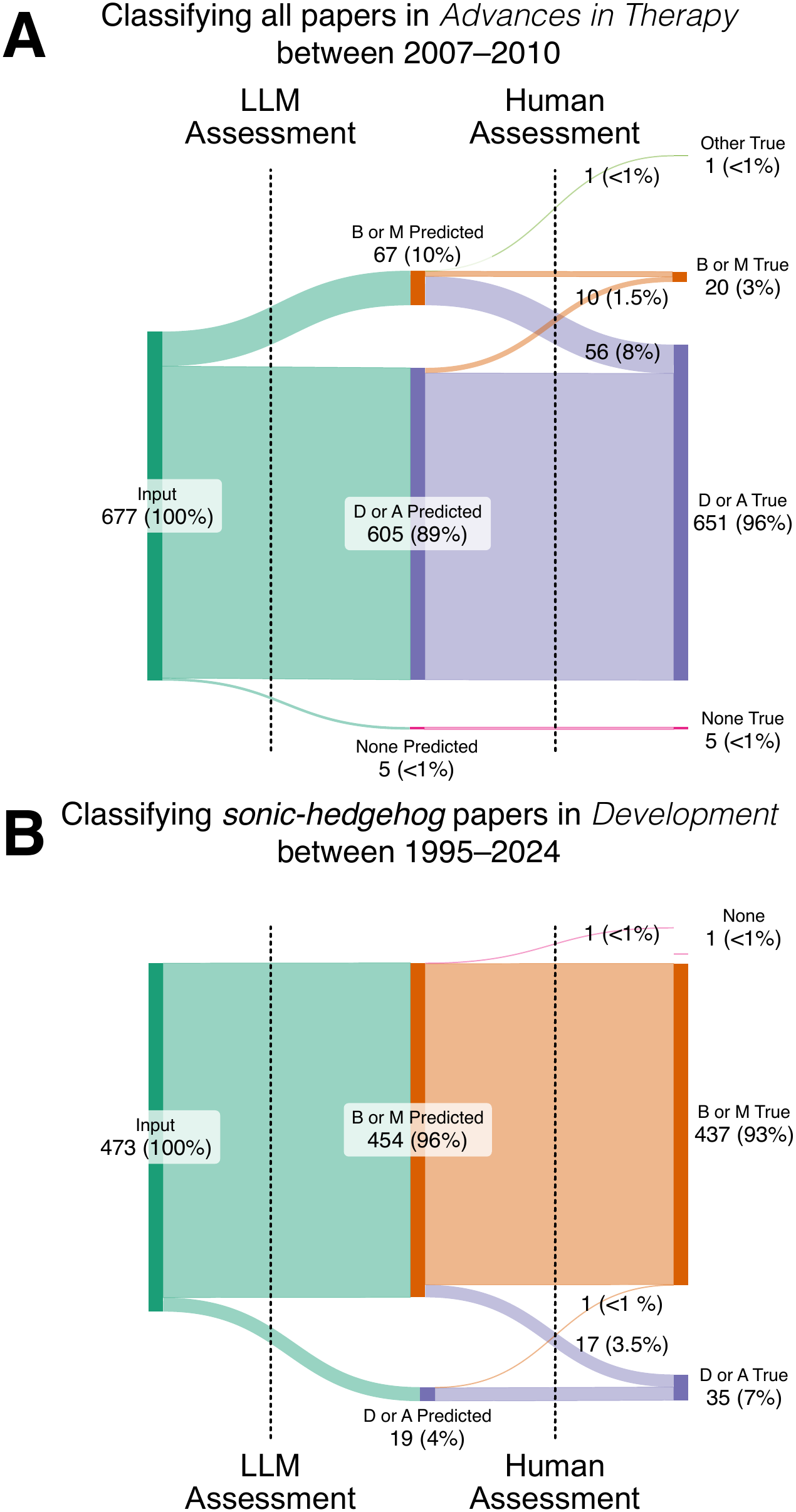
Test of accuracy of LLM. **A**. Classification of papers from *Advances in Therapy*. (left) 677 abstracts from the primarily disease and applied-focused journal *Advances in Therapy* were passed to the LLM. (center) The LLM predicted 605 were of the categories disease (D) or applied (A), 67 of the categories basic (B) or mechanistic (M), 5 had no abstract (None). (right) After human assessment of all abstracts, the true category of each abstract is shown. Misclassified abstracts are represented in the branches moving from one category to another after human assessment. The numbers on those branches show how many abstracts were misclassified: 56 abstracts classified as B or M were judged to better fit the category D or A, 10 abstracts classified as D or A were judged to be B or M, and 1 paper classified as B or M was not classifiable from the abstract. **B**. Same as above using papers published in *Development* between 1995–2024 that included the topic *shh* (*sonic hedgehog*).

**Table S1.** List of journals used for analysis, the category of journal, 5-year impact factor (IF) according to Web of Science Journal.

| **Category** | **High IF Journal** | **5-year IF** | **Low IF Journal** | **5-year IF** |
|---|---|---|---|---|
| *Multidisciplinary Sciences* | *Nature* | 55 | Proceedings of the Royal Society B-Biological Sciences | 4.4 |
|  | *Science* | 49.7 | Scientific Reports | 4.3 |
|  | *Nature Communications* | 17.2 | Journal of the Royal Society Interface | 3.8 |
|  | *Science Advances* | 14.1 | Heliyon | 3.9 |
|  | *PNAS* | 10.6 | Royal Society Open Science | 3.2 |
| *Biology, Cell Biology, Biochemistry & Molecular Biology* | *Cell* | 48.9 | International Journal of Molecular Sciences | 5.7 |
|  | *Molecular Cell* | 17.7 | Journal of Cell Science | 3.8 |
|  | *Developmental Cell* | 11.4 | Developmental Biology | 2.5 |
|  | *EMBO Journal* | 10.6 | FEBS Journal | 5.6 |
|  | *Current Biology* | 9.3 | BMC Biology | 5.4 |
|  | *Cell Reports* | 8.1 | Molecules | 5 |
|  | *PLOS Biology* | 7.7 | PLoS One | 3.2 |
|  | *Cell Metabolism* | 33.4 | Biofactors | 5.7 |
|  | *Cell Death & Differentiation* | 13.7 | Development Growth & Differentiation | 1.7 |
| *Genetics & Heredity* | *Nature Genetics* | 37.4 | Genetics | 4.1 |
|  | *Molecular Biology and Evolution* | 11.9 | Genome Biology and Evolution | 3.2 |
|  | *Genome Biology* | 16.3 | G3 Genes Genomes Genetics | 2.5 |
| *Biotechnology, Biochemical research methods* | *Nature Biotechnology* | 59.5 | BMC Biotechnology | 3.6 |
|  | *Nature Methods* | 51.7 | New Biotechnology | 5 |
| *Immunology* | *Immunity* | 33.2 | Frontiers in Immunology | 6.8 |
| *Neurosciences* | *Nature Neuroscience* | 24.8 | Frontiers in Cellular Neuroscience | 4.9 |
|  | *Neuron* | 16.6 | Journal of Neuroscience | 4 |
| *Ecology, Evolution* | *Nature Ecology & Evolution* | 17.6 | Evolution | 3.1 |
| *Microbiology* | *Cell Host & Microbe* | 20.5 | PLoS Pathogens | 5.4 |

**Table S2. Results of LLM Classifier for categorization of all high IF journal abstracts in our dataset. (available from** https://figshare.com/s/e959259f1f4bd22ad9be)

**Table S3. Results of LLM Classifier for categorization of abstracts from *Advances in Therapy* 2000–2010, including human assessment.**

**Table S4. Results of LLM Classifier for categorization of abstracts from *Development* 1995-2024 containing the keyword ‘shh’, including human assessment.**

**Table S5.** Correlations for the proportion of publications (in all selected journals) for each model organism along the years. *, p < 0.05; **, p < 0.01; ***p < 0.001; NS, non-significant.

| **Organism** | **Pearson correlation coefficient (1995 - 2004)** | **Pearson correlation coefficient (2005 - 2014)** | **Pearson correlation coefficient (2015 - 2024)** |
|---|---|---|---|
| **Mouse** | 0.90*** | -0.68* | -0.98*** |
| **zebrafish** | 0.96*** | -0.87*** | -0.95*** |
| ***Xenopus*** | -0.97*** | -0.98*** | -0.91*** |
| ***Drosophila*** | 0.12 (NS) | -0.99*** | -0.99*** |
| ***Arabidopsis*** | 0.97*** | -0.87** | -0.88*** |
| ***C. elegans*** | 0.95*** | -0.95*** | -0.96*** |
| **yeast** | 0.50 | -0.99*** | -0.98*** |
| ***E. coli*** | -0.98*** | -0.98*** | -0.98*** |

**Table S6.** Correlations for the proportion of publications (in all high IF journals) for each model organism along the years. *, p < 0.05; **, p < 0.01; ***p < 0.001; NS, non-significant.

| **Organism** | **Pearson correlation coefficient (1995 - 2004)** | **Pearson correlation coefficient (2005 - 2014)** | **Pearson correlation coefficient (2015 - 2024)** |
|---|---|---|---|
| **Mouse** | 0.73* | 0.68* | -0.93*** |
| **zebrafish** | 0.94*** | 0.73* | -0.89*** |
| ***Xenopus*** | -0.97*** | -0.97*** | -0.87** |
| ***Drosophila*** | 0.78** | -0.91*** | -0.95*** |
| ***Arabidopsis*** | 0.98*** | 0.13 (NS) | -0.85** |
| ***C. elegans*** | 0.95*** | -0.73* | -0.83** |
| **yeast** | -0.44 (NS) | -0.97*** | -0.93*** |
| ***E. coli*** | -0.97*** | -0.95*** | -0.88*** |

**Table S7.** Correlations for the proportion of publications (in all low IF journals) for each model organism along the years. *, p < 0.05; **, p < 0.01; ***p < 0.001; NS, non-significant.

| **Organism** | **Pearson correlation coefficient (1995 - 2004)** | **Pearson correlation coefficient (2005 - 2014)** | **Pearson correlation coefficient (2015 - 2024)** |
|---|---|---|---|
| **Mouse** | 0.95*** | -0.73* | -0.98*** |
| **zebrafish** | 0.89*** | -0.98*** | -0.93*** |
| ***Xenopus*** | -0.95*** | -0.97*** | -0.91*** |
| ***Drosophila*** | -0.83** | -0.99*** | -0.99*** |
| ***Arabidopsis*** | 0.77** | -0.63 (NS) | -0.83** |
| ***C. elegans*** | 0.67* | -0.92*** | -0.95*** |
| **yeast** | -0.37 (NS) | -0.97*** | -0.98*** |
| ***E. coli*** | -0.92*** | -0.04 (NS) | -0.99*** |

**Table S8.** Correlations for different conference attendances along the years. *, p < 0.05; **, p < 0.01; ***p < 0.001; NS, non-significant; NA, not available.

|  | **Total participants** | **Exhibitors/staff/speakers/comps** | **Faculty/lab technicians** | **Post-docs** | **Graduate students** | **Undergraduate students** | **Others** |
|---|---|---|---|---|---|---|---|
| ***C. elegans*** | -0.16 (NS) | 0.55 (NS) | 0.16 (NS) | -0.78* | -0.05 (NS) | 0.06 (NS) | -0.22 (NS) |
| **yeast** | -0.91 (NS) | 0.90 (NS) | -0.86 (NS) | -0.96* | -0.91 (NS) | 0.20 (NS) | NA |
| ***Drosophila (GSA)*** | 0.27 (NS) | 0.67 (NS) | -0.39 (NS) | -0.13 (NS) | 0.31 (NS) | 0.64 (NS) | 0.00 |
| ***Drosophila (EDRC)*** | 0.61 (NS) | NA |  |  |  |  |  |
| ***zebrafish*** | 0.85** | NA |  |  |  |  |  |

**Table S9.** Correlations for the counts of publications (in all selected journals) for each model organism along the years. *, p < 0.05; **, p < 0.01; ***p < 0.001; NS, non-significant.

| **Organism** | **Pearson correlation coefficient (1995 - 2004)** | **Pearson correlation coefficient (2005 - 2014)** | **Pearson correlation coefficient (2015 - 2024)** |
|---|---|---|---|
| **Mouse** | 0.99* | 0.95*** | 0.57 (NS) |
| **zebrafish** | 0.96*** | 0.95*** | 0.09 (NS) |
| ***Xenopus*** | -0.97*** | 0.10 (NS) | -0.78** |
| ***Drosophila*** | 0.85** | 0.94*** | -0.95*** |
| ***Arabidopsis*** | 0.98*** | 0.92*** | 0.48 (NS) |
| ***C. elegans*** | 0.97*** | 0.92*** | -0.58 (NS) |
| **yeast** | 0.30 (NS) | 0.87** | -0.92*** |
| ***E. coli*** | -0.77*** | 0.93*** | -0.318 (NS) |

**Table S10. Raw data used to generate graphs of all figures.**

